# Small molecule activator of phosphatase PP2A remodels scaffold PR65 structural dynamics to promote holoenzyme assembly

**DOI:** 10.1101/2025.11.26.690749

**Authors:** Sema Z. Yilmaz, Anupam Banerjee, Satyaki Saha, Michael Ohlmeyer, Reuven Gordon, Laura S. Itzhaki, Ivet Bahar, Mert Gur

## Abstract

Small molecule activators of protein phosphatase 2A (PP2A), hereafter SMAPs, have attracted substantial interest, for their potential to inhibit cancer cell proliferation by targeting PR65, the scaffold subunit of the PP2A heterotrimer. PR65 is a uniquely flexible and stable molecule composed of 15 tandem HEAT (Huntingtin, Elongation factor 3–PP2A–TOR1) repeats. We characterized the binding sites and interactions of two SMAPs ATUX-8385 and DT-061 with PR65 and evaluated effects on PR65 structural dynamics using docking and molecular dynamics simulations. We initiated SMAP-bound PR65 simulations starting from two binding sites: S1, determined by cryo-electron microscopy for DT-061 bound to PP2A, on the inner helices of the HEAT repeats 2 and 3 (2_i_ and 3_i_); and S2, predicted by docking of ATUX-8385 onto PR65, on 4_i_ and 5_i_ and outer helices 5_o_ and 6_o_ consistent with footprinting experiments. S2 proved to be a stable site for both SMAPs when initiating the simulations at S2. However, neither DT-061 nor ATUX-8385 demonstrated stable binding to S1. DT-061 rapidly dissociated from S1 to settle instead at a neighboring site S4 overlapping with our previously identified S3 for PR65 in extended form, suggesting that binding to S1 may be a 2-step process: an initial binding to PR65 alone, either to S3/S4 or S2, followed by movement to S3/S4, and then an induced relocation to S1 upon complexation with the regulatory and catalytic subunits. Targeted in silico mutagenesis showed that mutations at S2 and S4 destabilized SMAP binding to the PR65 (subunit). Heterotrimeric PP2A simulations showed that S3 and S4 binding were not persistent upon complexation. Together, these results corroborate our findings. Furthermore, preferentially stabilized a relatively extended PR65 conformation that would accommodate, if not promote, the assembly of the catalytic and regulatory subunits to prompt the activation of the trimeric phosphatase.

## INTRODUCTION

The preservation of cell signaling and homeostasis is vital for the proper operation of living organisms, with any disruptions in these functions potentially contributing to the onset of various health conditions. Signaling pathways and cellular homeostasis are controlled by a sophisticated regulatory relationship between kinases and phosphatases, with phosphorylation and dephosphorylation activities, respectively.^1^ Abnormal kinase activation or phosphatase inactivation can trigger pathologic hyperphosphorylation, a factor implicated in the development of cancer and neurodegenerative disorders.^2, 3^ Despite a significant focus on kinase inhibitors as drug targets for disease treatment, the strategy of modulating phosphatases’ activation remains less explored though potentially rewarding.^4-9^ One crucial class of phosphatases is the serine/threonine protein phosphatase 2A (PP2A) family, which plays a key role in maintaining cellular homeostasis.^10-12^ Due to its extensive regulatory function, PP2A, which is often found to be dysregulated in various human diseases, presents itself as an attractive candidate for therapeutic intervention across a spectrum of human diseases.^4-9,13^

PP2A enzymes are heterotrimers, comprised of a scaffold (A) subunit known as PR65, a catalytic (C) subunit, and a regulatory substrate-recruiting (B) subunit (**Figure 1**). There are over 40 different B subunits^2^, which selectively control the PP2A substrate and hence the specific activity of PP2A. The diverse array of B subunits allows PP2A to exert various levels of control over a majority of cellular signaling pathways, including c-Myc^14^, MAPK/ERK kinase (MEK)^15^, and mammalian target of rapamycin (mTOR)^16, 17^. PR65 and the catalytic subunits form the core of PP2A, with PR65 providing a flexible platform for the assembly of the heterotrimeric complexes.^18^ PR65 experiences the highest frequency of mutations, particularly along the binding interface of the B subunits, and many such mutations have been implicated in altering the composition of the PP2A holoenzyme.^19-22^

**Figure 1.**
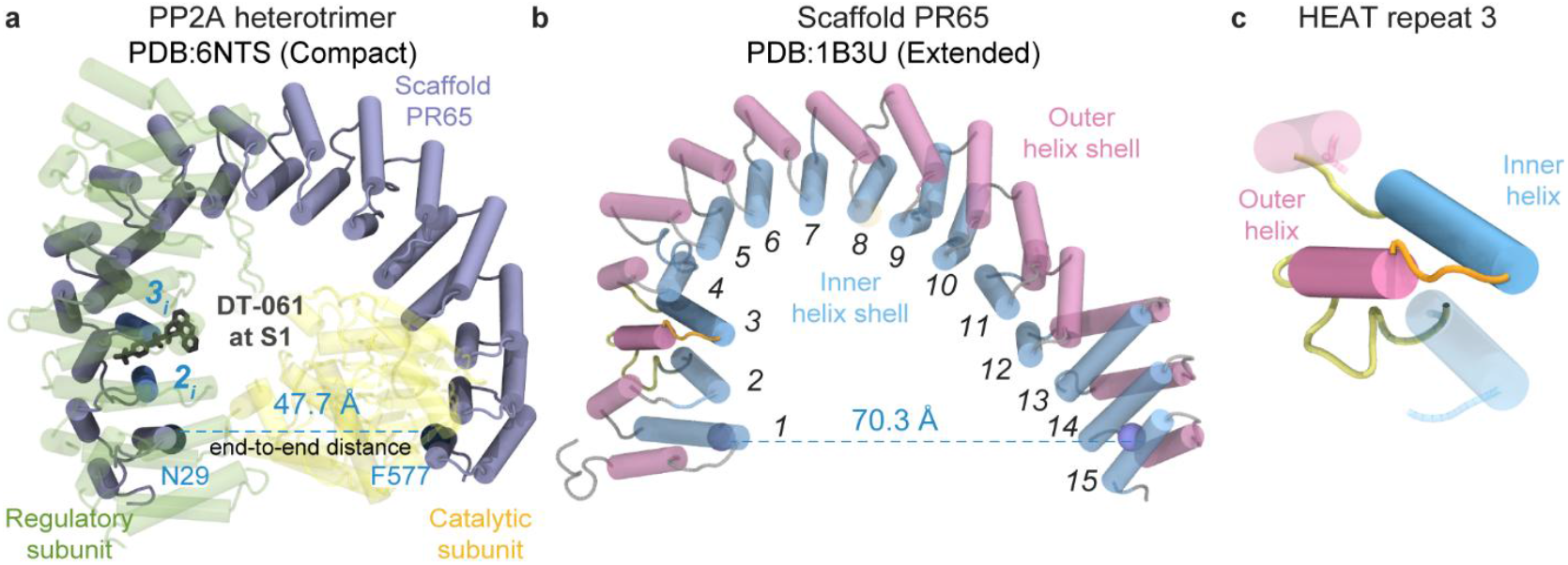
Schematic of the DT-061-bound heterotrimeric PP2A complex structure. (a) Cryo-electron microscopy (cryo-EM) structure of the PP2A heterotrimer bound to DT-061 (protein data bank (PDB): 6NTS^2^), shown in cartoon representation. PR65 (in compact conformation) is shown in *blue*; the catalytic and regulatory subunits are colored *yellow* and *green*, respectively. DT-061 is shown at binding site S1 (*black licorice*). (b) Extended PR65 structure (PDB: 1B3U^23^), with inner and outer shell helices of the 15 HEAT (Huntingtin, Elongation factor 3–PP2A–TOR1) repeats colored *blue* and *pink*, respectively. The degree of extension of PR65, or its end-to-end distance, is defined by residues N29 and F577 (shown as *beads*). The end-to-end distances are provided for both the compact (a) and extended (b) forms. (c) A HEAT repeat consists of two α-helices joined by a short interhelix segment. The neighboring inner and outer helices are shown in *semi-transparent cartoon representation*.

PR65 is a tandem repeat (TR) protein composed of 15 HEAT repeats, each consisting of approximately 40-residues folded in helical hairpins. Upon stacking in a one-dimensional fashion, these repeats give PR65 an extended, horseshoe-like super-helical structure composed of outer and inner helical shells. The architecture of PR65, together with other TR proteins containing ankyrin and tetratricopeptide motifs, is distinct from that of globular proteins. TR proteins, with their unique supersecondary structures, provide a highly flexible scaffold for molecular recognition, while their structural regularity also imparts high stability to enable them to function as scaffold.^24^ Their versatile nature allows them to adapt to various conformational changes based on their binding partners, functioning as adapter molecules or ‘hubs’. The regular repeat structure enables effective propagation of signals across the structure and plays a vital role in transmitting information within multi-subunit assemblies. As such, PR65 and similar scaffolding proteins composed of TRs are highly suitable for the dissection and redesign of their biophysical properties, which positions them as valuable targets in the fields of biotechnology and medicine.^18, 25-29^

Our previous work showed that the PR65 scaffold can fluctuate between extended and compact conformations, thus facilitating the binding and optimal interactions between the catalytic and regulatory subunits of PP2A.^18^ PR65 fluctuations may open or close the enzyme’s substrate binding/catalysis interface and alter the placements of specific catalytic residues to enable multiple cycles of substrate dephosphorylation.^30^ Understanding the impact of small molecule binding on PR65 structural dynamics is crucial in deciphering the molecular mechanisms underlying PP2A regulation. Our previous study^18^, using elastic network models (ENMs)^31^, molecular dynamics (MD) simulations, and hybrid methods combining the two, highlighted the role of PR65 in enabling PP2A enzymatic activity. The study underlined the remarkable flexibility of PR65, which accommodated changes in end-to-end distance of 20-40 Å between its compact and extended conformations. The study further highlighted the significant role of intra-repeat coils at the C-terminal arm of PR65 in allosterically mediating the collective dynamics of PP2A, thereby suggesting potential target sites for modifying PP2A function. In a more recent study^32^, we investigated the impact of single-point mutations on PR65 dynamics using a combination of *in situ* and *in silico* methods, including a total of 13.8 µs of all-atom MD simulations. Previous 2 ns-long steered MD (SMD) simulations^30^ to investigate PR65’s response to external uniaxial deformation provided insights into how the stretching and relaxation of the scaffold might facilitate the catalytic cycle.

Studies on the effect of small molecules binding on PR65 structural dynamics at the atomic level have been limited. In recent years, small molecule activators of PP2A (SMAPs), which are orally bioavailable compounds specifically designed to enhance PP2A activity, have attracted attention. SMAPs hold the potential to inhibit cancer cell proliferation by activating the PP2A.^33^ They were reported to operate by binding to the PR65 scaffold subunit and thereby driving conformational changes that affect PP2A.^33^ A high-resolution cryo-EM structure of PP2A in complex with the SMAP DT-061 showed that the DT-061 binds to a regulatory pocket lined by all three subunits of PP2A to selectively stabilize the PP2A-B56α holoenzyme^2^. PP2A-B56α complexation upon DT-061 binding induces cell death and reduces tumor growth through substrate dephosphorylation, including its well-characterized oncogenic substrate c-Myc.^2^ ATUX-8385 is another novel SMAP, and the ATUX-8385 bound PP2A structure is yet to be resolved. Specifically designed to activate PP2A without causing immunosuppression, ATUX-8385 demonstrated a decrease in hepatoblastoma proliferation, viability, and cancer cell stemness *in vitro*.^34^ We focus here on DT-061 as a structurally benchmarked ligand (cryo-EM resolved in the holoenzyme) and on ATUX-8385 as a water-soluble tricyclic sulfonamide tool compound that facilitates biophysical interrogation of PR65. The recently reported SMAP ATUX-1215^35^ retains the core diaryl–spacer–sulfonamide H-donor pharmacophore shared by DT-061 and ATUX-8385. A mechanistic understanding of how DT-061 or ATUX-8385 binding affects PR65 structure and dynamics is still lacking. Resolving such mechanisms of PP2A activation by SMAPs will pave the way for engineering novel molecules and broadening their therapeutic applications.

In this study, we explored the binding mechanisms of ATUX-8385 and DT-061 and determined their effect on the structure and dynamics of PR65. We employed a range of *in silico* techniques, including docking simulations and MD simulations (~25.4 µs cumulative duration). Our simulations of monomeric PR65 revealed a new site, called site S2, which exhibited a high affinity to bind SMAPs when PR65 was in its compact form. This site differed from that (site S1, between the inner helices of the HEAT repeats 2 and 3, shortly referred to as 2_i_ and 3_i_) observed in the cryo-EM structure resolved for DT-061 bound PP2A (PDB: 6NTS^2^). Once bound to S2, the SMAPs remained bound for extended durations (up to 700 ns) in multiple runs and exerted specific effects on PR65 conformational dynamics. In the monomeric PR65 simulations, DT-061 also populated a site on the N-terminal halves of 5_i_–7_i_ (S4), whereas the extended form favored^36^ the adjacent and partially overlapping site located near the inner helices 3_i_ and 4_i_ (S3). S3/S4 is proposed to be the region that first binds SMAP in the extended form of PR65, before assembly of the PP2A trimer during which the SMAP relocates by induced fit to the adjacent site S1. Targeted alanine and glutamic acid mutagenesis at S2 (Y154, R166, F191, and N199) and S4 (L221, and R257), followed by MD simulations of monomeric PR65 in complex with SMAPs confirmed these as hotspot and anchoring residues critical for SMAP binding. Complementary PP2A holoenzyme simulations further revealed that S3 and S4 are not persistent ligand-binding sites once the catalytic and regulatory subunits are assembled (i.e., when PR65 forms the trimeric PP2A complex), supporting a hierarchical relocation pathway. The newly identified binding sites and mechanisms, and their effects on PR65 dynamics provide new hypotheses for effective design and development of PP2A activators.

## RESULTS AND DISCUSSION

### Unbiased docking simulations reveal the dependency of the preferred binding sites of ATUX-8385 on PR65 conformation, extended or compact

As mentioned above, DT-061 interacts with the inner helices 2_i_ and 3_i_ in the heterotrimeric structure resolved by cryo-EM for DT-061-bound PP2A (PDB: 6NTS^2^). This experimentally resolved site is referred to as the PR65 site S1 (**Figure 1a**). There is no structural data on the binding of the SMAP ATUX-8385 to PP2A. Previously, experiments showed that radiolabeled DT-061 bound to the isolated PR65 subunit with K_D_ ≈ 235 nM, indicating that PR65 alone can engage DT-061.^33^ Furthermore, in the same study, hydroxyl radical footprinting experiments identified K193, E196, and L197 as likely coordinating residues at a potential SMAP-binding site for DT-061. Yet, the mechanism of SMAP binding or the sequence of events remains to be elucidated, e.g. whether PR65 presents a high-affinity site for the SMAP, the binding of which facilitates subsequent assembly of the heterotrimer, or whether the PR65 conformational state, open/extended or closed/compact, affects the binding site.

To address this knowledge gap, we first conducted blind (unbiased) docking simulations of ATUX-8385 onto PR65. We generated 266 possible binding poses for the compact (PDB: 6NTS^2^, chain A) and extended (PDB: 1B3U^23^) conformations of PR65 (**Figure 2a, c**; *red sticks*). Simulations showed that the high-affinity sites depended on the conformational state of PR65. In the compact/closed state, ATUX-8385 preferentially bound to the vicinity of HEAT repeat 5 (specifically outer helices 5_o_ and 6_o_), which also comprises the K193–L197 stretch (enclosed in *magenta sphere* in **Figure 2a**, and *magenta dot* in **Figure 2b**), consistent with the residues (K193, E196, and L197) inferred from hydroxyl radical footprinting experiments^33^. We also observed another high-affinity site between 3_i-_ 5_i_(including T97–V102) in the close neighborhood of site S1 (enclosed in *black circle;* **Figure 2a**; and *black dot* in **Figure 2b**). In the extended/open state of PR65, on the other hand, ATUX-8385 preferentially bound to the 3_i_-5_i_ near the S1, while also showing affinity to other sites on PR65 (**Figure 2c-d**). Notably, these respective sites overlap with the regulatory and catalytic subunits’ binding regions, pointing to an avidity for ligand binding to those sites upon the extension of the PR65 structure. To gain a clearer understanding of the docking conformations, specifically whether they reside at the trimeric protein interface or are solvent-exposed, we superposed the binding poses onto the trimeric PP2A structure (PDB: 6NTS^2^), in which the PR65 scaffold subunit adopts a compact conformation, as illustrated in **Figure S1**.

**Figure 2.**
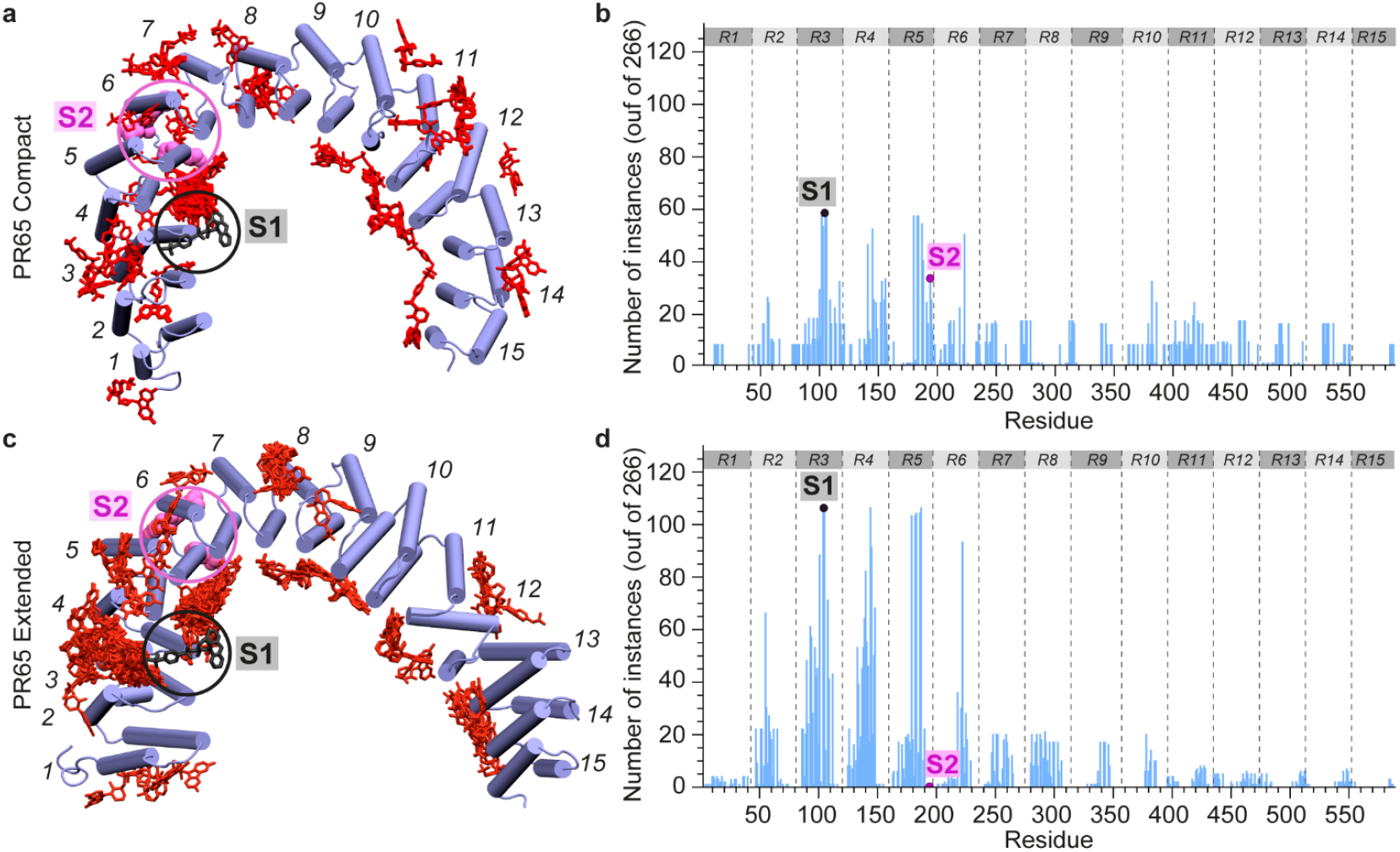
Results from blind docking simulations of SMAP ATUX-8385 to PR65 in compact and extended conformations. Results (266 docked poses) are shown from multiple docking simulations of ATUX-8385 (*red sticks*) onto PR65 (*blue cartoon*) in the compact (a-b) and extended (PDB: 1B3U^23^) forms (c-d). The distributions of the binding poses over the 15 HEAT repeats (denoted by *gray bars*, R1–R15, and separated by *gray dashed lines*) are shown in (b) and (d). The *blue bars* show the number of ATUX-8385 instances within 5 Å of each PR65 residue. The catalytic and regulatory subunits were not included in the simulations. Docked conformations in (a) were generated using blind docking via SwissDock and CB-Dock, while in (c) they were generated using blind docking via AutoDock Vina. DT-061 resolved by cryo-EM (PDB: 6NTS^2^) is shown in *black sticks* (inside a *black circle*) to indicate its binding site, S1. The *magenta spheres* (inside the *magenta circles*) refer to three residues, K193, E196, and L197, proposed to be involved in binding a SMAP to the PR65 scaffold.

The binding poses were further refined by guiding docking simulations for ATUX-8385 onto the compact PR65 using the K193-L197 residues as a constraint in the Rosetta Ligand Docking Protocol^37^ on the ROSIE server^38^, which led to a refined binding pose near E196 (**Figure S2b**)^39^. The latter, referred to as site S2, comprises the helices 4_i_, 5_i_, 5_o_, and 6_o_, and exhibited a binding energy of -10.5 kcal/mol, as calculated using PRODIGY-LIG_40_. Refinement of ATUX-8385 binding to the extended form, on the other hand, using Autodock Vina^41^ with K193–L197 site as a reference, uncovered another novel binding site (S3) located at 3_i_, 4_i_, and 5_i_ with a PRODIGY-LIG_40_ predicted affinity of -9.5 kcal/mol (**Figure S2c**).

Given the high sensitivity of SMAP binding to the conformational state of PR65, we next proceeded to the examination of the stability of the SMAPs ATUX-8385 and DT-061 when bound to the S1 and S2 sites, anticipating that a time-resolved analysis would provide more rigorous information beyond that inferred from docking simulations.

### MD simulations showed the high affinity of site S2 to stably bind SMAPs

We have carried out five sets of MD simulations (**Table 1**), with each set comprising three 704 ns long runs. Sets I-V were performed using as starting systems the apo form of PR65 (set I), and PR65 in the presence of DT-061 bound to S1 (set II), ATUX-8385 bound to S2 (set III), DT-061 bound to S2 (set IV), and ATUX-8385 bound to S1 (set V). **Figure S2a-b** illustrates the initial positions adopted for each of the SMAPs bound to the two sites, as well as the initial conformers (compact) of PR65 (**see Methods**).

**Table 1.** Simulated WT monomeric PR65 systems and their durations. (*)

| Set of runs | Starting binding state | Observed SMAP binding site | Trajectory length for observed SMAP binding (ns) |  |
| --- | --- | --- | --- | --- |
|  |  |  | Individual runs | Combined |
| I | Apo | Apo | (a) 704, (b) 704, (c) 704 | 2112 |
| II | DT-061 at S1 | S4 | (a) 692, (b) 560, (c) 88 | 1340 |
| III | ATUX-8385 at S2 | S2 | (a) 704, (b) 704, (c) 704 | 2112 |
| IV | DT-061 at S2 | S2 | (a) 704, (b) 704, (c) 704 | 2112 |
| V | ATUX-8385 at S1 | S5 | (a) 614, (b) 0, (c) 0 | 614 |
|  |  | S6 | (a) 0, (b) 600, (c) 0 | 600 |

(\*) All runs were conducted in triplicate, each of duration 704 ns. Columns 4 and 5 refer to the period of time the SMAPs were observed to be bound to the site indicated in column 3, except for the first row (without a SMAP) where the durations of the simulations are listed.

Simulations initiated with either SMAP bound to S2 indicated stable binding in all triplicate runs (sets III a-c and IV a-c), clearly demonstrating the strong affinity of SMAPs to bind to site S2. Principal component analysis (PCA)_42-44_ of SMAP conformations sampled in the combined trajectories revealed that ATUX-8385 may assume three slightly different conformations (designated as S2-A1, S2-A2, and S2-A3), while stably binding to the same site S2 (**Figure 3a**); DT-061 also steadily remained bound to the same site, with a minor conformational rearrangement (S2-D in **Figure 3b**). The two panels in **Figure 3c** clearly show that the same site is selected with minor conformational changes that accommodate the specific SMAPs. **Figure 3d** provides a detailed description of the SMAP binding poses and local interactions at site S2.

**Figure 3.**
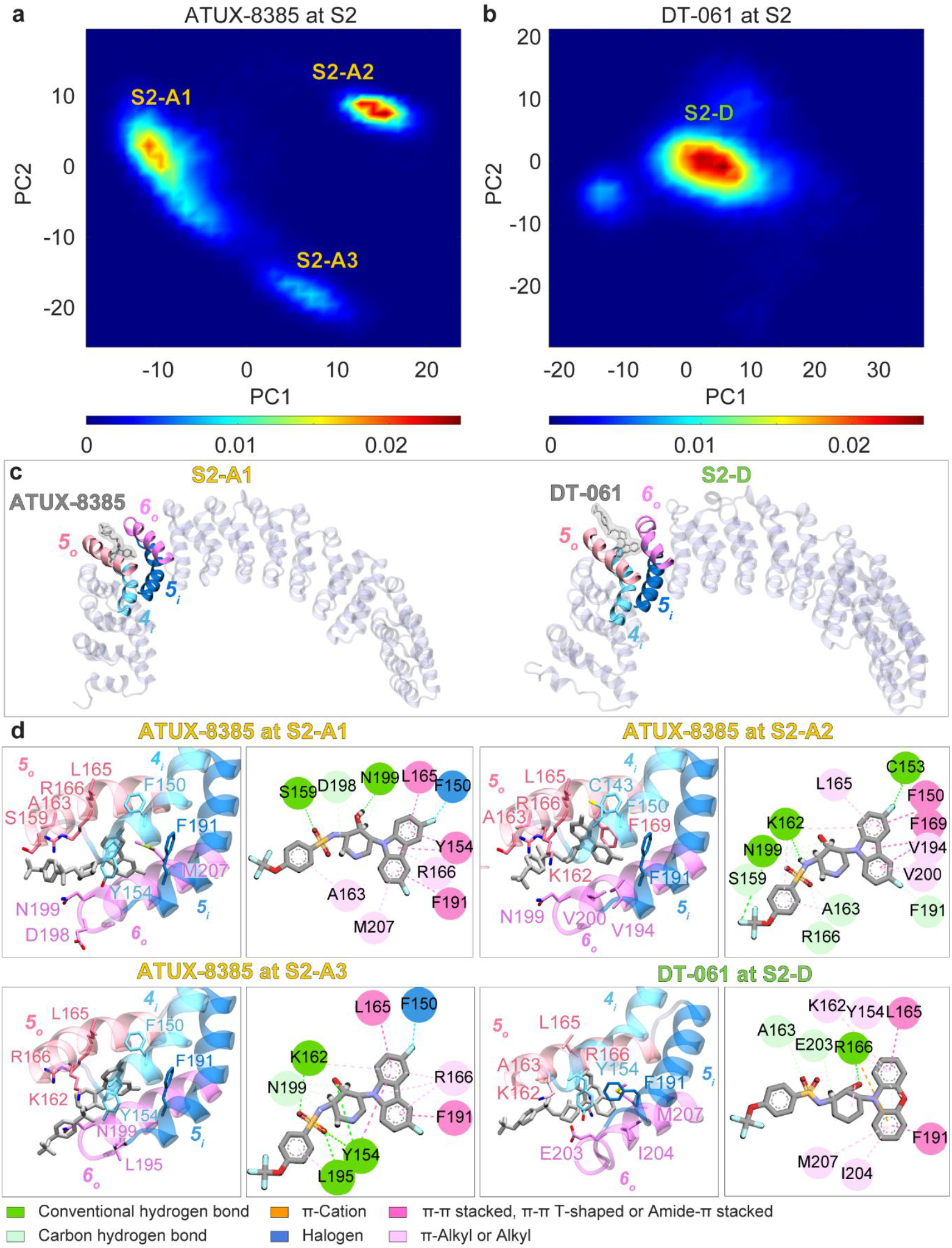
Extensive MD simulations demonstrate the high stability of SMAPs bound to site S2. (a-b) Distributions of SMAP binding conformations projected onto the two-dimensional reduced spaces defined by the first two principal components (PC1 and PC2) extracted from the respective MD trajectory sets III and IV (**Table 1**) showed that the same site may accommodate multiple conformations, S2-A1, S2-A2, and S2-A3 for ATUX-8385, and S2-D for DT-061, all bound to the same site S2. (c) The overall view of PR65 complexed with ATUX-8385 (*left*), and DT-061 (*right*), both bound to S2. (d) The detailed local interactions and binding poses of the SMAPs at each of the identified substates at site S2, as described by the labels. Each substate is depicted through two distinct representations: (*left*) a diagram generated using VMD^**45**^, that displays SMAPs and surrounding residues in a licorice representation, and (*right*) a schematic illustration, from the Discovery Studio Visualizer ^**46**^, which showcases the atomic interactions. The types of interactions are indicated by the color code (described in the inset).

During the three runs conducted for ATUX-8385 bound to S2 (set III a-c), the conformation S2-A1 was most frequently sampled with a cumulative time of 1328 ns, whereas S2-A2 and S2-A3 were observed for 481 ns and 295 ns, respectively. The S2-D conformation of DT-061 was observed for most of the 2112 ns of simulations (**Table 1**). The root-mean-square deviation (RMSD) between representative S2-A1 and S2-A2 conformations of ATUX-8385 was 4.6 Å upon aligning C_α_ of the PR65 helices forming the binding site, and that between S2-A1 and S2-A3 was 4.7 Å. Collectively, these changes in SMAP bound pose and conformations while remaining bound to PR65 highlight the accessibility of different ATUX-8385 orientations inside the binding pocket S2 and the favorable entropic contribution to binding.

ATUX-8385 made extensive interactions with PR65 4_i_ residues F150 and Y154, 5_o_ K162, L165, R166, 5_i_ F191, and 6_o_ E196, and N199 (**Figures 3d**, and **S3-S5**). In addition to these shared interactions between the conformations S2-A1, S2-A2, and S2-A3, S2-A1 exhibited interactions with 5_o_ S159, and A163, and 6_o_ V194, L195, D198, V200, E203, I204, and M207, while S2-A2 formed additional interactions with 4_i_ S151 and C153, 5_o_ S159, A163, and F169, 5_i_ E190, and 6_o_ V194 and V200. S2-A3 formed additional interaction with 5_o_ R170, 6_o_ L195. The most frequently observed interactions in S2-A1 were the hydrogen bonds/ hydrophobic interactions with Y154 (π-π T shaped/π-π stacked), K162 (π-alkyl), and R166 (amide-π stacked/π-alkyl), hydrogen bonds with S159, D198, and N199, hydrophobic interactions with A163 (π-alkyl) and F191 (π-π T shaped/π-π stacked), and π-sulfur/hydrophobic interaction (π-alkyl) with M207. Those of S2-A2 were the hydrophobic interactions with F150 (π-π T-shaped), Y154 (π-π T-shaped), L165 (π-alkyl), F169 (π-π T-shaped), and V194 (π-alkyl), hydrogen bond/halogen interaction with C153, and hydrogen bond/hydrophobic interaction (π-alkyl) with K162, and hydrogen bonds with S159, A163, R166, F191, and N199. In S2-A3, the most frequent interactions were the halogen interaction with F150, hydrogen bonds/hydrophobic interactions with Y154 (π-π T shaped), and L195 (π-alkyl), hydrophobic interactions with L165 (amide-π stacked) and R166 (π-alkyl/π-σ), and hydrogen bonds with K162 and N199.

In the binding pose S2-D, DT-061 exhibited interactions with 4_i_ residue Y154, 5_o_ residues S159, K162, A163, L165, R166, and F169, 5_i_ residue F191, 6_o_ residues V200, E203, I204, and M207 (**Figures 3d**, and **S6**). The most frequently observed interactions were the hydrophobic interactions with Y154 (π-π T shaped/π-alkyl), L165 (amide-π stacked), F191 (amide-π stacked/π-π stacked), V200 (π-alkyl), and M207 (π-alkyl), hydrogen bond with S159, and hydrogen bonds/hydrophobic interactions with A163 (π-alkyl), and R166 (amide-π stacked/π-alkyl).

### DT-061 exhibits a strong tendency to dislocate from S1 and settle in a new site, S4

In contrast to their observed high affinity for site S2, both SMAPs exhibited rapid dissociation from S1 in sets II and V. This observation is consistent with the cryo-EM structure of PP2A in complex with DT-061 (PDB: 6NTS^2^), which reveals that S1 becomes an intersubunit pocket only when PR65 is complexed with the catalytic and regulatory subunits at the intersubunit interface. DT-061 dissociated within 1 ns (**Movie S1**); and re-attached through its phenoxazine group to a new binding site on the N-terminal halves of 5_i_-7_i_, designated as S4 (**Figures 4 and S7**). The identification of this alternative site took between 12 to 145 ns (**Figures 4 and S7**). More than half of the trajectories generated in set II showed DT-061 bound to S4, highlighting the high stability of SMAP DT-061 bound to that site. **Figure 4c** illustrates three snapshots sampled by the SMAP during one of the trajectories, and the final stable conformation (*rightmost diagram*) stabilized at that site. Similar poses observed in the other two trajectories are presented in **Figure S7**, corroborating the high affinity of S4 for binding DT-061. The main interaction partners of DT-061 at S4 were R182 of 5_i_, D217, L221, L222 of 6_i_ and W256, R257, and Y260 of 7_i_ (**Figures 4c, S7**, and **S8**). Among these interactions, those most frequently observed were hydrophobic interactions (π-alkyl) with L221 and hydrogen bond/electrostatic interaction (π-cation)/halogen interaction with R257. R257 and L221 acted as anchors for the attachment of phenoxazine, enabling DT-061 to explore various conformations at site S4.

**Figure 4.**
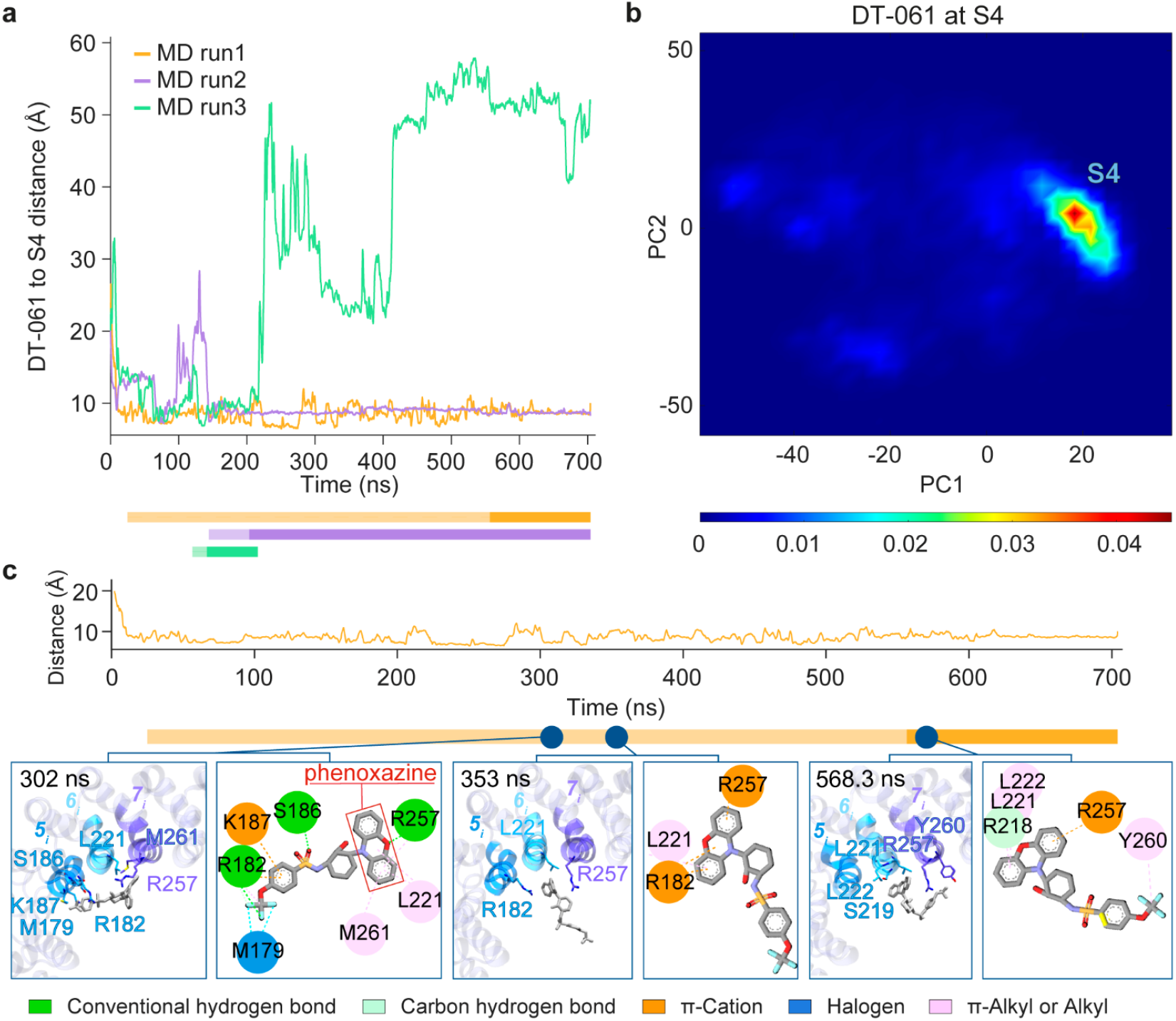
A relocation of DT-061 from S1 to the novel binding site S4 is consistently observed in three independent MD runs. (a) The evolution of distances between DT-061 and S4 in simulations initiated with DT-061 at S1. The distance was defined as the length of the vector between DT-061’s center of mass (CoM) and the CoM of the N-terminal halves of segments 5_i_-7_i_ (P178-A183, D217-L222, and R257-M261). Bars under the figures indicate the trajectory stretches that were used for PCA. The *darker portions* of the bars represent the parts of the trajectory that predominantly sample the minimum in the principal space-projected map shown in (b). (b) SMAP binding poses projected onto the principal space of bound conformers extracted from MD trajectories. (c) Closeup view of one of the trajectories showing the preferential binding site throughout most of the trajectory, and the most stable binding pose attained at the end of the run, also stabilized for extended durations in a 2^nd^ run (shown in *purple* in (a)) and a short duration in the 3^rd^ (*green*, time interval corresponding to the *darker green bar* section). See the results for the other two runs in **Figure S7**, which confirms the preferential binding to S4.

As to ATUX-8385, it also underwent a quick (<1 ns) dislocation from S1 (**Movies S2-S3**), and it failed to identify a consistent site in the three runs: in one run it completely dissociated from PR65; in the 2^nd^, it found a new site at 8_o_ and 6_i_-8_i_; and in the 3^rd^ it bound another site at 12_o_, and 12_i_-14_i_ (**Figure S9**). These relatively short-lived binding poses, respectively designated as S5 and S6 were sampled for total durations of 614 ns and 600 ns (**Table 1**), indicating the lower stability compared to DT-061 bound to S4. See **Figure S10** for more details on these short-lived sites and comparison with the pose S2 preferred by ATUX-8385.

Comparison of this S4 site with the above mentioned site S3 (**Figure S2c**), also reported in our recent work^39^ revealed several overlapping residues among those coordinating the two SMAPs, including R182, A183, S186, and K187. Moreover, after 800 ns of MD simulation with ATUX-8385 bound at S3, the overlap between the S3 and S4 sites increased to include residues R182, A183, S186, K187, and L222 (**Figure 5**). We performed additional docking simulations of DT-061 onto the extended conformation of PR65 using Autodock Vina^41^, employing residues K193– L197 as a reference. These simulations further confirmed binding to S3 with the involvement of residues such as D105, V108, R112, T144, A183, and K187 that coordinate either SMAP, ATUX-8385 or DT-061 (**Figure S2c**), and the overlaps with site S4 A183 and K187. The docked poses of DT-061 and ATUX-8385 at S3 exhibited a binding affinity of –9.5 kcal/mol, as calculated by PRODIGY-LIG^40^.

**Figure 5.**
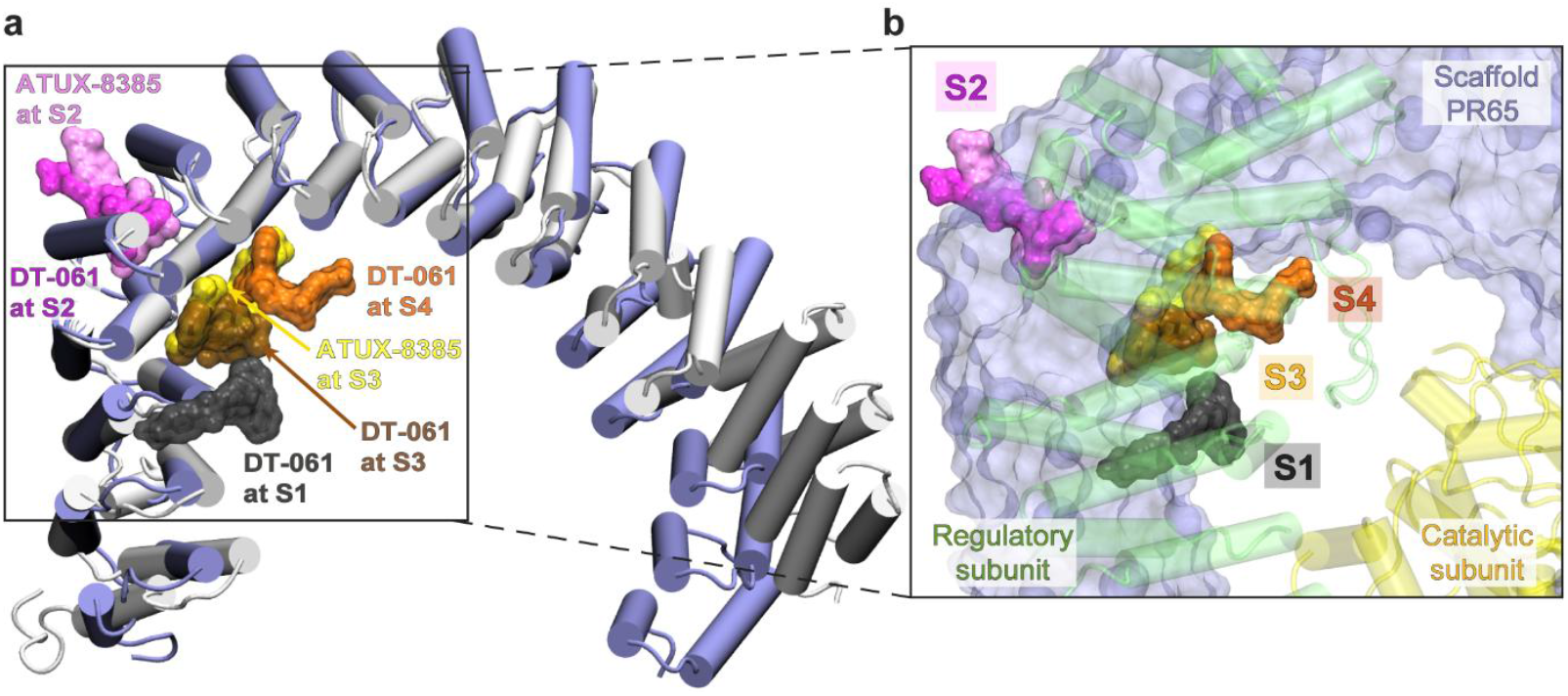
MD explored S4 site is in close proximity to S1 and previously identified site S3. (a) Structural alignment of compact (*blue cartoon*) and extended (*gray cartoon*) conformations of PR65. Representative SMAP binding poses at sites S1-S4 are depicted in surface representation. All binding poses, except DT-061 at S1 (cryo-EM) and S3 (docked), are MD-refined binding poses. (b) A zoomed-in view of the binding region, with PR65 rendered in *surface representation*. The regulatory and the catalytic subunits are shown as *transparent green* and *yellow cartoons*, respectively.

Therefore, we suggest that site S4 may represent part of an extended binding region that encompasses S2, S3, and S4. Furthermore, the data support a sequential binding model in which SMAPs engage the S3/S4 region either after an initial encounter at S2 or by direct binding, and in both cases ultimately relocate to S1 upon complexation with the regulatory and catalytic subunits.

### Targeted *in silico* mutagenesis confirms the role of S2 and S4 residues for SMAP binding

To assess the role of residues within the S2 and S4 sites for SMAP binding, we performed targeted *in silico* mutagenesis at S2 and S4. For S2, we introduced alanine and glutamic acid substitutions at Y154, R166, F191, and N199 (**Table 2**). In the MD simulations (set VI; ~500 ns each) of the alanine-substituted systems (Y154A, R166A, F191A, and N199A), ATUX-8385 dissociated within ~250 ns in one MD run and adopted a shifted pose in the other two, whereas DT-061 rotated rapidly in three MD runs (set VII) and failed to return to its native orientation **(Figures 6 and S11)**. Glutamic acid substitutions (Y154E, R166E, F191E, and N199E) allowed ATUX-8385 to remain bound in shifted poses compared to WT (set VIII). By comparison, DT-061 rotated within the S2 site in all simulations of the glutamic acid mutant (set IX, ~400 ns each). These findings confirm that this cluster of residues forms as hotspots that stabilize SMAP binding at S2.

**Table 2.** Simulated PR65 mutant systems and their durations.

| Set of runs | Mutations | Starting conformation | Observed binding change | Simulation duration (ns) |
| --- | --- | --- | --- | --- |
| VI | Y154A, R166A, F191A, and N199A | ATUX-8385 at S2 (Starting conformation of set III) | (a) Left from S2, (b-c) Shifted binding pose at S2 | (a) 504, (b) 504, (c) 504 |
| VII | Y154A, R166A, F191A, and N199A | DT-061 at S2 (Starting conformation of set IV) | (a-c) Rotated binding pose at S2 | (a) 504, (b) 504, (c) 504 |
| VIII | Y154E, R166E, F191E, and N199E | ATUX-8385 at S2 (Starting conformation of set III) | (a-c) Shifted binding poses at S2 | (a) 404, (b) 404, (c) 404 |
| IX | Y154E, R166E, F191E, and N199E | DT-061 at S2 (Starting conformation of set IV) | (a-c) Rotated binding poses at S2 | (a) 404, (b) 404, (c) 404 |
| X | L221A, and R257A | DT-061 at S4 (MD-refined pose of set II b, taken at 577.7 ns) | (a) Left from S4, (b-c) Rotated binding poses at S4 | (a) 504, (b) 504, (c) 504 |
| XI | L221E, and R257E | DT-061 at S4 (MD-refined pose of set II b, 577.7 ns) | (a-c) Left from S4 | (a) 404, (b) 404, (c) 404 |

**Figure 6.**
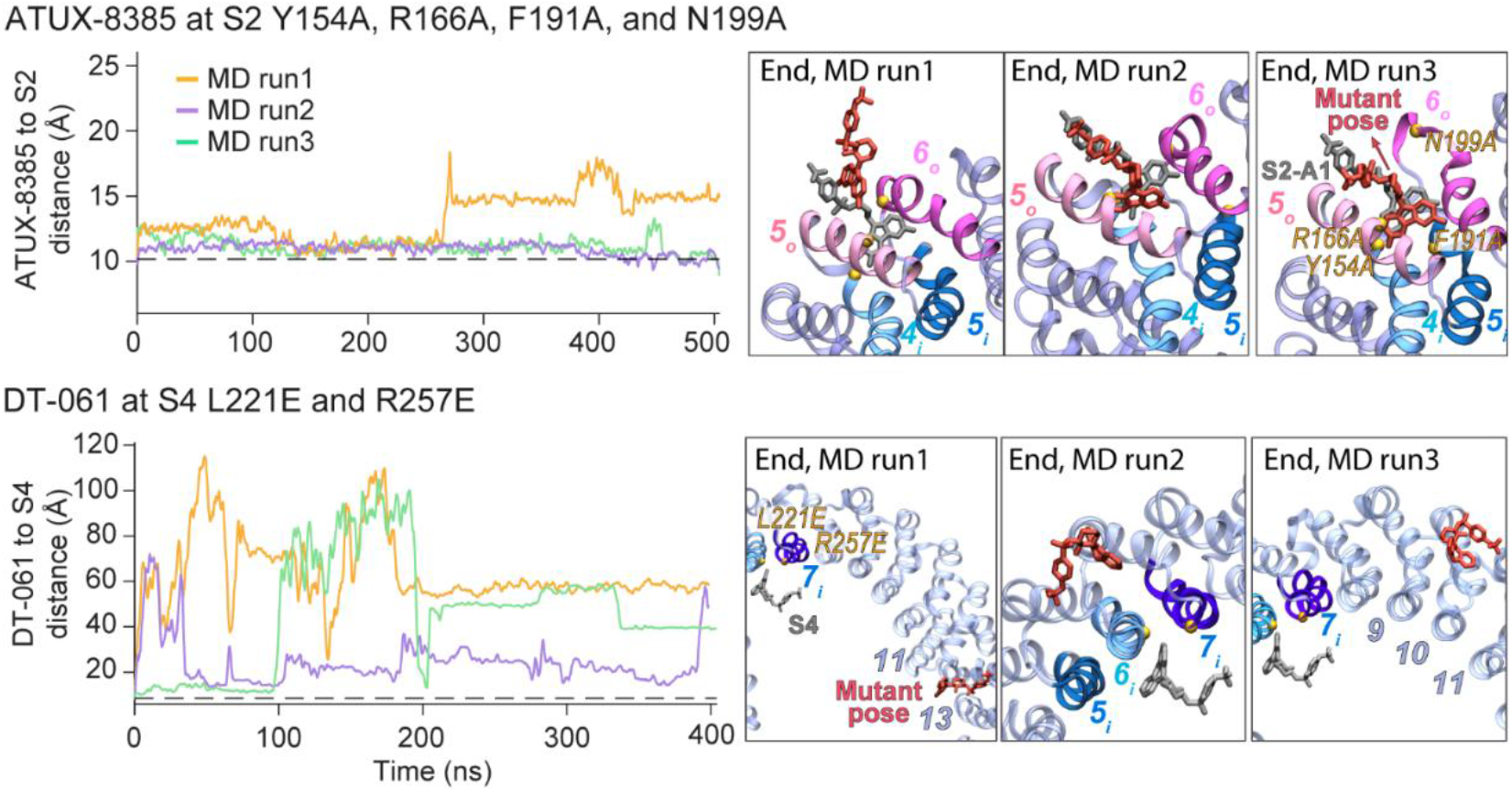
Time evolution of SMAP–binding site distances in mutant MD simulations and endpoint poses. Left panels: For simulations initiated with SMAPs bound at S2, the distance of each SMAP to the S2 site was calculated as the separation between the CoM of each SMAP and the CoM of PR65 helices 4_i_, 5_i_, 5_o_, and 6_o_ (residues F140–Y154, A160–S174, P178–A192, and L197–S213, respectively). For simulation initiated with SMAPs at S4, the distance between the SMAP and S4 was calculated as described in **Figure 4.** Each trace (*orange, purple*, and *green*; run1-3, respectively) represents one of three independent MD runs starting from the same binding pose and PR65 mutant. Right panels: Endpoint snapshots showing the final bound pose captured at the last frame of each trajectory for ATUX-8385 or DT-061 in the indicated mutant backgrounds.

For S4, we performed *in silico* mutagenesis of residues L221 and R257, again introducing both alanine and glutamic acid substitutions (**Table 2**). In the alanine-substituted systems (L221A and R257A), DT-061 remained bound in most runs (set X, each of ~500 ns length) but dissociated in one run, suggesting reduced stability compared to the WT. By comparison, glutamic acid substitutions (L221E and R257E, set XI, ~400 ns each) consistently led to unbinding of DT-061 across all replicates, confirming that the electrostatic repulsion introduced by these mutations disrupted ligand anchoring (**Figures 6 and S11**). Together, these results establish L221 and R257 as primary anchoring residues that stabilize DT-061 at S4.

### S3 and S4 are not persistent ligand sites in the trimer

To test whether the single-subunit pockets preferred on PR65 alone (S3/S4) destabilize once the holoenzyme is assembled and further support the suggested S3/S4 → S1 transition of SMAPs upon complexation with the regulatory and catalytic subunits, we seeded those poses into the PP2A trimer and ran unbiased MD simulations (**Table 3**). First, the MD-refined S4 pose of DT-061 identified on PR65 was superposed onto the trimer and simulated (set XII, three runs, ~500 ns each). DT-061 detached from S4 in two of three runs, progressing toward S1 in one run, while in the other it explored the PR65–B56α interface without forming a persistent binding pose. (**Figures 7** and **S12a**). We then positioned DT-061 at S3 by placing the docked DT-061 pose from the PR65 system (**Figure S2c**) into the trimer (set XIII, three × ~300 ns runs, **Figure S12b**). DT-061 left S3 within ~10 ns in one run, for the remaining simulation time it lingered on the inner helix shell of PR65 without forming a well-defined, long-lived pose and instead made only intermittent contacts with PR65; in the other two runs, it remained bound at S3 (**Figure S12b**).

**Table 3.** Simulated PP2A heterotrimer systems and their durations.

| Set of runs | Starting binding state | Observed binding change | Simulation duration (ns) |
| --- | --- | --- | --- |
| XII | DT-061 at S4 (MD-refined pose of set II b, 577.7 ns) | (a-b) Left from S4, (c) Stayed at S4 | (a) 504, (b) 504, (c) 504 |
| XIII | DT-061 at S3 (Docked pose) | (a-b) Stayed at S3, (c) Left from S3 | (a) 304, (b) 304, (c) 304 |
| XIV | ATUX-8385 at S3 (Docked pose) | (a-b) Left from S3, (c) Stayed at S3 | (a) 304, (b) 304, (c) 304 |
| XV | ATUX-8385 at S3 (MD-refined end pose of MD simulations initiated from ATUX-8385 at the S3) | (a-c) Left from S3 | (a) 304, (b) 304, (c) 304 |

**Figure 7.**
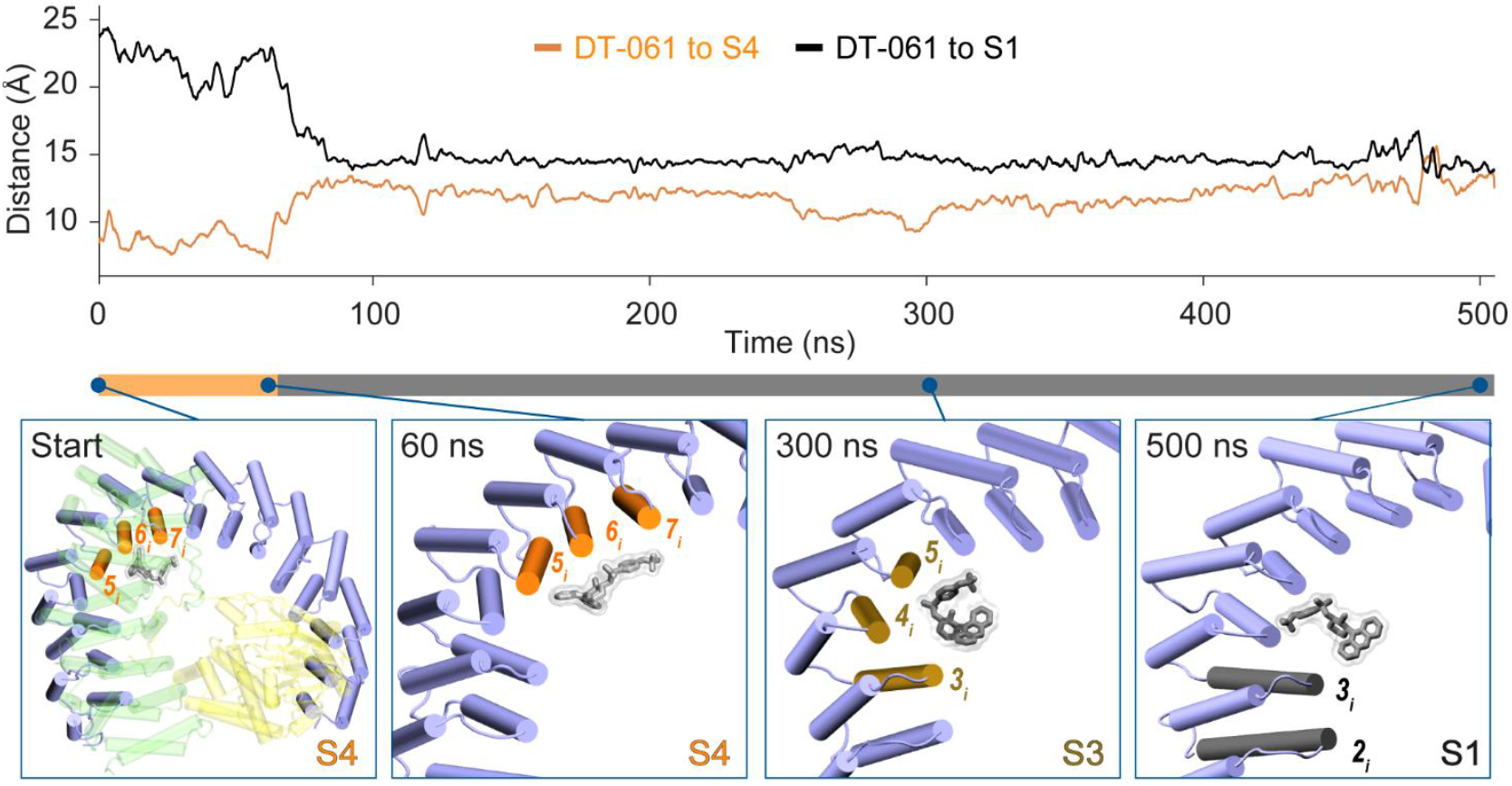
DT-061 seeded at S4 migrates toward S1. *Top:* Time evolution of CoM distances between DT-061 and site-defining segments. The S4 reference (*orange*) is the CoM of the N-terminal halves of PR65 helices 5ᵢ–7ᵢ; the S1 reference (*black*) is the CoM of the N-terminal halves of helices 2ᵢ–3ᵢ (residues D62–A67 and T101–A107), representing subsets of helices that form part of the S4 and S1 sites, respectively. The *horizontal bar* below the plot indicates the approximate position of DT-061 over the course of the simulation, *orange* when DT-061 resides at S4 and *gray* after it departs from S4. *Bottom:* Representative snapshots illustrating the trajectory of DT-061 relative to binding sites. S4, S3, and S1 are highlighted in *orange, brown*, and *black*, respectively. The *bottom-left panel* shows the starting configuration with DT-061 in S4; subsequent panels (60 ns, 300 ns, and 500 ns) depict its progression from S4 to S3 and ultimately to S1.

We performed an analogous test with ATUX-8385. Three MD runs (each ~300 ns) were performed for each of the following starting models: ATUX-8385 was added to the trimer in (i) its S3 docking pose on PR65 (set XIV) (**Figure S2c**) and (ii) its MD-refined S3 pose from the PR65-only system (set XV) (**Table 3**). ATUX-8385 dissociated from S3 in five of these six runs, and as was the case for DT-061, sampled the PR65–B56α interface rather than S3/S4 (**Figure S13**). Taken together, these holoenzyme simulations show that neither DT-061 nor ATUX-8385 is stably retained at S3 or S4 once the regulatory and catalytic subunits are present.

### Multiplicity of high affinity sites near repeats 5-6 suggests a 2-step process consisting of binding of DT-061 to S4 or S3, followed by induced fit to S1 in favor of PP2A heterotrimer formation

Given the multiplicity of sites observed in docking and MD simulations, and in experiments, we sought to identify an overarching property. We noticed that SMAPs localized near the same region yet exhibited distinct binding-site dislocations specific to each SMAP type and PR65 conformation.

In all three MD simulations initiated with DT-061 bound at the S1 site of the PR65, rapid DT-061 dissociation was consistently observed. This behavior, at odds with the binding site S1 observed for DT-061 by cryo-EM resolution of the ternary complex PP2A, suggests that the observed site S1 is stably formed only upon the complexation of PR65 with the other two subunit(s) of PP2A. Interestingly, following dissociation from S1, DT-061 exhibited spontaneous re-association with the nearby S4 site in all MD trajectories.

Although the S1 site is not stable in monomeric PR65, a broad region in its vicinity presents multiple adjacent pockets identified by docking simulations (S2 and S3) and confirmed (S2 and S3) or refined (S4) by MD simulations. This region consistently emerges as a high avidity location for SMAP initial attachment to the PR65 scaffold and later resettling to optimize the interactions with the catalytic and regulatory subunits (**Figure 5**). The trimer PP2A simulations further substantiate this mechanism by showing that S3/S4 do not constitute long-lived binding modes once the regulatory and catalytic subunits are present.

In the trimeric PP2A structure (PDB: 6NTS^2^), DT-061 at S1 interacts with B56α residues I237, Y238, K283, K316, and F317, in addition to residues on PR65. However, we observed a more extensive interaction network for DT-061 at S2 when bound to monomeric PR65 compared to this interaction network between DT-061 and B56α in the PP2A trimer. This further supports that initial capture by PR65 is the more probable binding route, rather than a hypothetical alternative pathway involving primary binding to B56α.

Taken together, our findings suggest a sequence of events in which DT-061 initially associates with the monomeric PR65. It binds either the S4/S3 sites or the S2 site depending on PR65 conformational state, and from S2 may relocate to the adjacent S4 and S3 sites, the latter presenting a high affinity in the extended form. Binding of DT-061 stabilizes the extended form that facilitates the recruitment of other PP2A subunits, which subsequently prompts an induced fit ultimately stabilizing the SMAPs at the S1 site (S3/S4 → S1). The proximity of S4, S3, and S1 further supports this translocation hypothesis. However, the functional relevance and dynamics of S3 warrant more extensive investigations. As to ATUX-8385, the S2 site predicted by docking simulations, and confirmed by MD, further emphasizes that the conformational adaptability of the PR65 repeats 5 shared with S3 and S4 plays an important role in PP2A function. As illustrated in **Figures S3-S6 and S8**, the SMAPs might assume an ensemble of conformations at sites S2 and S4, which presumably adds to the stability of the SMAPs at those sites. A closer look at the effect of the conformational (structure and dynamics) change in PR65 is presented next to understand how these small molecules act as activators.

### SMAP binding impacts the conformational state and dynamics of the scaffold

We compared the equilibrium dynamics of PR65 in the apo state to those in the presence of SMAPs bound to different sites (S2–S6). **Figure 8a** displays the root-mean-square fluctuation (RMSF) profile of amino acids for different cases; and **Figure 8b** illustrates the extreme conformations (*left*, compact; and *right*, extended) as well as an average conformation (*middle*) sampled by PR65 in apo form, and bound to ATUX-8385 to S2, to DT-061 at S2, to DT-061 at S4, (*from top to bottom*).

**Figure 8.**
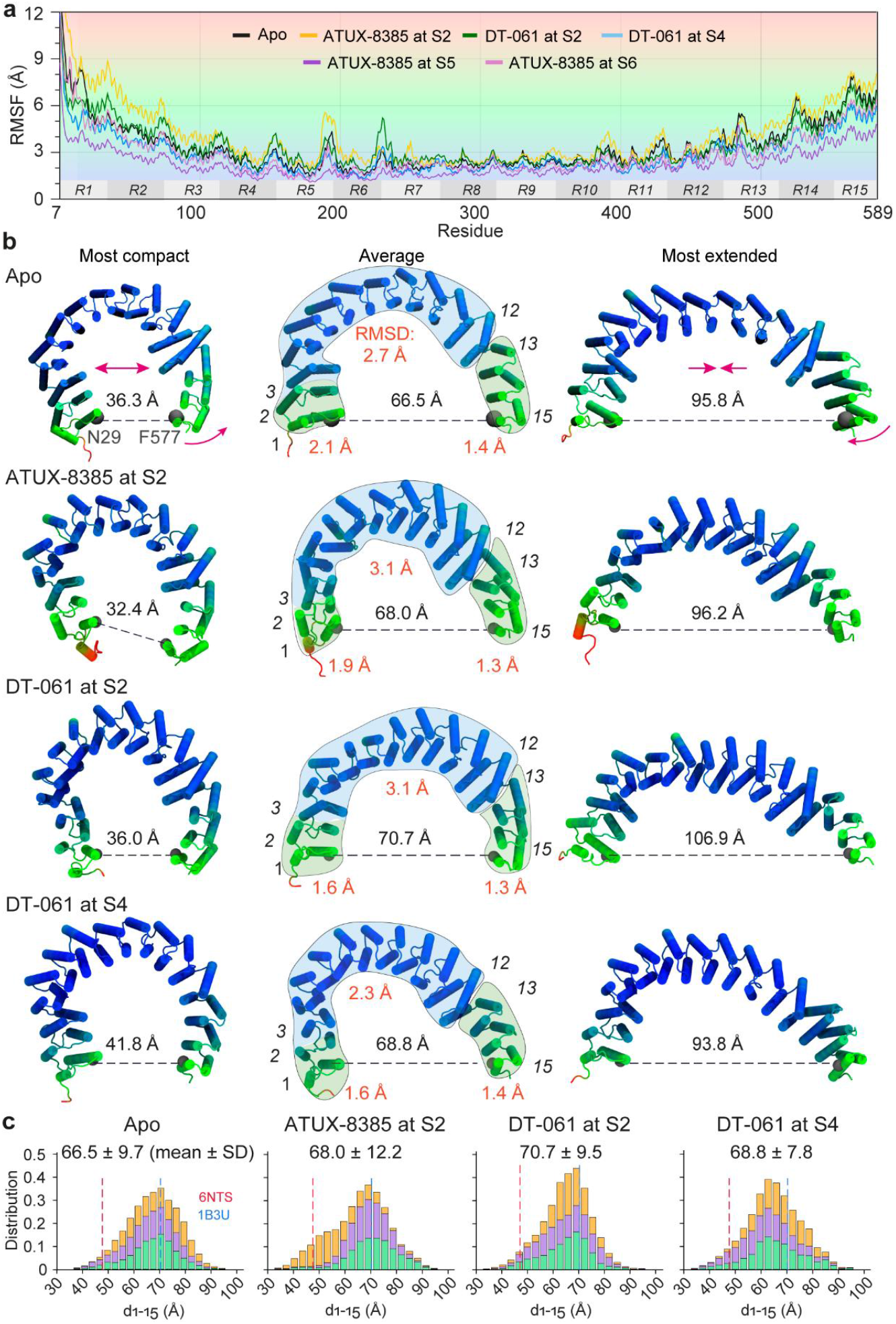
Residue mobilities observed for apo PR65 and its states bound SMAPs. (a) RMSF profiles of PR65 residues observed in MD simulations for the apo PR65 and five SMAP-bound states. The *gray boxes* labeled R1–R15 denote the corresponding HEAT repeat segments of PR65 along the sequence. (b) Conformations sampled during simulations. Each row illustrates the most compact (*left*), average (*middle*), and most extended (*right*) forms. The residues are colored in accordance with their RMSFs, as delineated in panel (a). The end-to-end distances are shown. In the *middle* structures, semi-transparent bubble overlays highlight the N-terminal (*green*), central (*blue*), and C-terminal (*green*) regions, with the mean internal RMSD values for each region annotated in *red*. The C- and N-termini show relatively small RMSDs, while the middle segment undergoes large internal rearrangements. All RMSD calculations were performed using C^α^ only. Rigid-like reorientation at the interface between repeats 12 and 13 further assists in the changes in end-to-end distance. (c) Histograms of end-to-end distances observed in the four types of simulations; each composed of three runs. *Dashed vertical lines* depict the end-to-end distances observed in the compact (PDB: 6NTS^2^) and extended (PDB: 1B3U^23^) structures of PR65.

The simulations demonstrated that the conformational fluctuations of apo PR65 permit sampling of the end-to-end distances observed in the compact (47.7 Å, trimeric) and extended (70.3 Å, monomeric) experimental structures, with a mean end-to-end distance of 66.5 Å averaged over three runs (**Figure 8c and S14)**. The difference between the end-to-end distance apo PR65 subunit observed in simulations and experiments is likely due to the non-physiological conditions under which the apo PR65 crystal structure (PDB: 1B3U^23^) was obtained (100 K, pH 5.5, lattice contacts). This issue was emphasized in our earlier study showing that crystallography and MD probe proteins under fundamentally different thermodynamic environments that can lead to structural differences^47^, as well as differences in the intrinsic dynamics of the protein. Within the MD simulations, the SMAP-bound average PR65 conformations are all more extended than the average apo PR65 MD conformations, by 1.5 Å to 4.2 Å. Among all MD runs, the most extended PR65 conformation was observed for DT-061 at S2 with an end-to-end distance of 106.9 Å.

Thus, SMAP binding induces an extension in PR65, which, in principle, could facilitate the accommodation of the catalytic and regulatory subunits (note that after complexation with these two subunits PP2A reverts to a compact form). This is consistent with the principle that ligand-induced changes exploit pre-existing soft modes of the scaffold, as shown by Meireles et al. (2011)^48^, with SMAP binding biasing PR65 toward an extended intermediate that facilitates assembly before compaction. The extension originates mainly from a cooperative rearrangement/ expansion in the middle portion (colored *blue*) as well as a hinge site between repeats 12 and 13 which enables the rotational motion of the C-terminal portion.

Histograms of PR65 end-to-end distances, measured as the instantaneous distance between N29 and F577 C^α^ atoms (representative of the N- and C-termini) are depicted in **Figure 8c** for each run, also showing the tendency to stabilize the extended conformer; and more information on the time evolution of end-to-end distances for each run is presented in **Figure S14a**, while **Figure S14b** shows their distributions as violin plots. Notable, a wider distribution (12.2 Å standard deviation, SD) is observed when ATUX-8385 is bound to S2, compared to both apo (9.7 Å SD) and DT-061 at S4 (7.8 Å SD). Further investigation showed an interesting behavior in that case. In two of the runs, a highly extended form (> 73 Å) of PR65 was stabilized, while in the third, a compact form dominated the large part of simulations, which later transitioned into an extended form. Consequently, a notably larger mean C^α^ atom RMSF was observed for ATUX-8385 at S2 (4.1 Å) compared to all other SMAP bound PR65s (**Figure 8a**). This was followed by apo and DT-061 at S2 (both 3.4 Å). All other SMAP-bound PR65s exhibited lower average RMSF than that of apo PR65 (2.2-2.9 Å).

As a final test, we performed a PCA^42-44^ of the SMAP-bound PR65 MD trajectories to identify the ability of the different structures to undergo global opening/closing motion essential to PP2A activity. For this analysis, only the coordinates of PR65 were used, excluding those of the SMAPs. We then assessed the similarity between these PCs and the deformation vector, which represents the overall change (3N-dimensional vector, for N C^α^ atoms) from the compact to the extended PR65 structure, by calculating the correlation cosines between them. Our PCA analysis (**Figure S14c**) shows that compact ↔ extended transitions are encoded in PR65’s intrinsic motions, and SMAP binding modulates the extent and propensity of these transitions.

By definition, PC1 describes the softest, and therefore energetically most favorable, collective mode of the scaffold. In accordance with the end-to-end distance characteristics which demonstrated that ATUX-8385 at the S2 site undergoes fewer transitions between the compact and extended forms, the combined trajectory exhibited a lower correlation cosine (of 0.60) for PC1 (**Figure S14c**); but this decrease was offset by an increase in PC2’s capability, which improved to 58%. Overall, the top four PCs yielded cumulative correlation coefficients comparable to the apo, suggesting that the adaptability of the structure to functional changes as driven by the softest modes was not reduced by SMAP binding, while its predisposition to binding the regulatory and catalytic subunits was enhanced by favoring more extended forms of the scaffold.

## CONCLUSIONS

This study provides new insights into the binding mechanisms of SMAPs ATUX-8385 and DT-061 to the PP2A scaffold PR65, and into their effects on the structure and dynamics of PR65. We identified a high affinity site, designated S2, for ATUX-8385 through docking simulations. MD simulations in triplicate confirmed the stability of this site, offering atomic-scale details into the interactions and conformational dynamics of ATUX-8385 at that binding pocket. Simulations also showed that the S2 site can accommodate three distinct binding conformations of ATUX-8385 in addition to the binding of DT-061, endowed by the conformational adaptability of that specific region (centered around HEAT repeat 5, and the surrounding helices, mainly 4_i_, 5_i_, 5_o_, and 6_o_). This conformational adaptability further points to the entropic contribution to stabilizing the bound state.

MD simulations of PR65 complexed with DT-061 bound to S1, on the other hand, revealed that this binding site identified by cryo-EM study of the PP2A heterotrimer, was not a high-affinity binding site for PR65 alone. Similarly, ATUX-8385 originally bound to S1 also dissociated within a short time from the scaffold PR65. This observation, seemingly at odds with the cryo-EM data, suggests that S1 is not necessarily a good binding site for SMAPs when PR65 is not complexed with the other subunit, but is likely to become so upon binding of the two additional (catalytic and regulatory) subunits, which indeed coordinate DT-061 in the cryo-EM structure. In accord with this inference, simulations of DT-061 binding to PR65 pointed to another binding site, S4, in close vicinity, which DT-061 consistently located after rapid dissociation from S1 in three independent runs. This recurrent behavior suggests that S4 serves as a first recognition site and allows for relocation of DT-061 to the nearby binding site S1 upon trimerization. S4 lies directly adjacent to, and partially overlaps with S3, the site we previously identified for ATUX-8385 by docking on the extended PR65 subunit. MD simulations now confirm S3 and its partial overlap with S4 for both SMAPs.

The Boltzmann averaged binding free energies calculated via PRODY-LIG^40^ were -9.6 kcal/mol for ATUX-8385 at S2, and -9.3 kcal/mol for DT-061 at S2, -8.1 kcal/mol for DT-061 at S4. These results suggest that SMAP ATUX-8385 exhibits the strongest affinity for S2 when PR65 scaffold protein is unbound to the other two subunits; this is followed by DT-061. These data highlight the high affinity of the S2 site to bind SMAPs. Yet S4 is selected in both compact and extended forms of SMAP, pointing to a favorable entropic contribution. Together, these results define S2–S4 as a contiguous binding region, arising from its solvent-exposed topology and residue composition enriched in aromatic and basic side chains, that captures SMAPs through favorable hydrophobic interactions and hydrogen bonds, accompanied by ligand-based entropic stabilization.

Collectively, our data supports the hypothesis of a potential hierarchical binding mechanism. According to this proposed mechanism, SMAPs engage as a first step either the high-affinity S2 pocket consistent with footprinting experiments or the sites S3 and S4 strongly suggested by MD simulations. Notably, S3 and S4 are both coordinated by 5_i_; S3 further shares 4_i_ and 5_i_ with S2, and 3_i_ with S1. So, conceivably, this site may serve as a bridge for the translocation of the SMAP to its cryo-EM resolved pose in the PP2A trimer, induced upon assembly of the three subunits of PP2A. As SMAP binding to PR65 scaffold promotes a conformational extension without affecting the scaffold flexibility or dynamics, the SMAP-bound PP2A shows a higher propensity (than unbound PP2A) to bind the catalytic and regulatory subunits, before the SMAP finally repositions into the heterotrimer-specific S1 pocket. This sequential pathway (S3/S4 → S1 or potentially S2 → S3/S4 → S1) provides a plausible mechanism linking ligand migration on PR65 to the stepwise assembly of the PP2A holoenzyme.

Our targeted *in silico* mutagenesis further supports this mechanism by independently confirming the importance of hotspot residues in stabilizing SMAP binding. Quadruple substitutions at Y154, R166, F191, and N199 at S2 weakened or destabilized SMAP binding. Similarly, double substitutions at L221 and R257 at S4 disrupted DT-061 binding, particularly with glutamic acid mutations, highlighting their anchoring role. These mutagenesis results provide an additional layer of validation for our predicted binding sites and demonstrate how computational approaches can guide experimental mutagenesis in future work.

Furthermore, alignment of PP2A holoenzymes containing distinct B family members [B56α (PDB: 6NTS^2^), B56γ (PDB: 2NPP^49^), B56δ (PDB: 8U1X^50^), B56ε (PDB: 8UWB^51^), B55α (PDB: 3DW8^52^), and PR70 (PDB: 4I5N^53^)] revealed that the extended SMAP-binding region in PR65 (sites S1–S4) is structurally conserved, with helix-only RMSD values of 0.8–1.2 Å relative to the 6NTS^2^ structure (DT-061 bound, B56α), while overall PR65 helical RMSD values ranged from 0.9– 4.4 Å (**Figure S15**). This conservation suggests that SMAP-binding hotspots are structurally preserved across different B subunits.

Finally, both DT-061 and ATUX-8385 influence PR65 structure and dynamics, promoting an extended form predisposed to supporting PP2A heterotrimer formation and thereby PP2A activation. Such conformational changes likely facilitate assembly of multiple B subunit PP2A heterotrimers, including non-classical RNAPII–Integrator– AC complexes that promote transcriptional termination^54^. Dissecting the effects on cellular signaling with different activator chemotypes remains an active area of investigation. Our findings offer a deeper mechanistic understanding of the role of SMAPs in modulating the activity of PP2A, which could be exploited for novel therapeutic interventions.

We note that PR65 extension alone does not necessarily linearly predict activity: our prior data with the non-functional SMAP DBK-776^36^ indicate that excessive extension (~80 Å) may actually impair activity, whereas ~70 Å appears to be closer to optimal. We therefore conclude that activity emerges from a balance between the degree of extension, modulation of intrinsic fluctuations, other ligand-induced conformational changes upon trimer assembly. While limited by the number of SMAPs studied here, our results suggest an avenue for future investigations into how distinct ligands remodel PR65 to tune holoenzyme assembly and function.

## METHODS

This work is purely computational; no compounds were synthesized or used experimentally. For reference, synthetic routes to the constrained tricyclic sulfonamide SMAP chemotype modeled here (e.g., DT-061 and ATUX-8385) are disclosed in Ohlmeyer & Kastrinsky, US Patent 9,937,180 B2 (2018)^55^, and Ohlmeyer & Zaware, US Patent 10,759,790 B2 (2020)^56^.

### Modeling of apo and SMAP-bound PR65 structures

The DT-061 bound trimeric PP2A structure (PDB: 6NTS^2^) determined with a resolution of 3.63 Å was used to model the apo structure of monomeric PR65 in a compact form. The coordinates for DT-061 were taken from the same structure to generate a DT-061 bound form of PR65 at the S1 site. ATUX-8385 structural model was generated using OpenBabel^57^. The coordinates for ATUX-8385 docked at the S2 site were obtained from our docking simulations, as described in the **Results and Discussion** section. To model the binding of ATUX-8385 at the S1 site, we aligned ATUX-8385’s carbon atoms (C11-C17 and C19-C22) against those of DT-061 (C1-C3, C6-C12, and C18) as shown in **Figure S2a**. For modeling DT-061 binding at the S2 site, we reversed this alignment procedure, aligning DT-061’s carbon atoms against those of ATUX-8385 at S2 (**Figure S2b**).

To evaluate the effects of targeted substitutions on SMAP binding at S2 and S4, PR65 mutants bearing the binding-site mutations described in the **Results and Discussion** section were constructed using the Mutator plugin in VMD^45^. The DT-061–bound conformation at the S4 site (from MD simulation set II b, 577.7 ns time instant) and the DT-061– and ATUX-8385–bound conformations at the S2 site (starting conformations of sets IV and III, respectively) were used as templates for mutation.

### Modeling of SMAP-bound PP2A structures

To obtain the apo PP2A template, the bound DT-061 ligand was removed from the trimeric PP2A structure (PDB: 6NTS^2^). Missing residues R295–L309 in the catalytic subunit were taken from the PP2A holoenzyme (PDB: 2IAE^58^) after rigid-body superposition of C^α^ atoms K4–R294 of the catalytic subunits (resulting C^α^ RMSD 0.7 Å); the C-terminal L309 was modeled in its methylated form, and the two Mn^2+^ ions resolved in 6NTS^2^ were retained. For SMAP-seeded trimer models, ligand poses determined on monomeric PR65 (see **Results and Discussion** section) were ported into the trimer by rigid-body alignment of PR65 C^α^ atoms N29–N229 between the PR65-only model and the 6NTS^2^ scaffold, followed by placement of the ligand in the aligned coordinates.

### MD simulations

Each apo and SMAP-bound structure was solvated in a water box containing explicit TIP3P water molecules, with 144 Å edge length in all directions. To neutralize the system and set the ion concentration to 150 mM NaCl, Na^+^ and Cl^-^ ions were added. The system sizes were approximately 286,900 and 283.900 atoms for PR65 and PP2A systems, respectively. All system preparation steps were conducted using VMD^45^.

MD simulations were conducted using the NAMD 3 software^59^, with the CHARMM36 all-atom force field^60^. SMAPs were parameterized using CHARMM-GUI Ligand Reader module^61^. A 2 fs time step was used. The temperature was maintained at 310 K via Langevin dynamics, utilizing a damping coefficient of 1 ps^−1^, and the pressure was held at 1 atm using the Langevin Nosé–Hoover method with an oscillation period of 100 fs and a damping time scale of 50 fs. Van der Waals interactions were calculated with a cut-off distance of 12 Å, and the particle-mesh Ewald method was applied for long-range electrostatic interactions.

Two rounds of system minimization and equilibration were executed before each production run. Initially, the protein structure was kept fixed and subjected to 10,000 minimization steps, followed by a 1 ns of equilibration. This first round of minimization-equilibration was designed to equilibrate the solvent around the protein. Subsequently, we performed a second round of minimization-equilibration, in which the system underwent an additional 10,000 step minimization without any restrictions on protein structure and dynamics, followed by a 2 ns of equilibration, applying harmonic restraints (k=1 kcal/mol/Å^2^) only on the C^α^ atoms. After these preparatory simulations, we removed all restraints and initiated production runs.

Results from 15 sets of MD simulations using NAMD 3 software^59^ (sets I-XV) are presented. Of these, 13 (sets II, IV, and V-XV) are newly performed, and two (sets I and III) were performed earlier and partially reported in the context of a comparison with experiments^32, 39^. The in-depth analysis of these two sets, as presented here, has not been previously performed. In addition to these 15 sets, we conducted^36^ an additional set of MD simulations initiated from the ATUX-8385 docked pose at the S3 site (**Figure S2c**), utilizing AMBER20 software^62^ with different simulation parameters and force fields. For details of the additional set of MD simulations please refer to **Supplementary Methods**.

### PCA

Details on PCA calculations can be found in previous studies.^42-44^ SMAP conformations were sampled with a frequency of 0.1 ns from MD trajectories. We aligned the SMAP-bound PR65 conformations via the interacting PR65 helices and tracked the relative movements of SMAPs. All calculations were conducted using our custom analysis codes, which were run in VMD and MATLAB. These codes also incorporated some of the built-in functions of these platforms. PCA systematically breaks down the movements seen in the trajectories into 3N-6 components, where N is the number of atoms used in the PCA calculations. Therefore, the first component (PC1) portrays the most dominant global changes in conformation, while PC2 characterizes the second most dominant movement. Distributions obtained by projecting SMAP non-hydrogen atom coordinates onto the first two principal components (PCs 1-2) showed the existence three main binding poses for ATUX-8385 at S2 (S2-A1, S2-A2, and S2-A3), one for DT-061 at S2 (S2-D) (**Figure 3**) in addition to one for DT-061 at S4 (**Figure 4**), one for ATUX-8385 at S5, and one for ATUX-8385 at S6 (**Figure S10**).

## ASSOCIATED CONTENT

### Supporting Information

Supplementary methods describing the MD simulations for ATUX-8385 bound to PR65 at S3, and supplementary figures S1–S15 (PDF)

Supplementary movie S1 (AVI)

Supplementary movie S2 (AVI)

Supplementary movie S3 (AVI)

## AUTHOR INFORMATION

### Authors

Sema Z. Yilmaz − Department of Computational and Systems Biology, School of Medicine, University of Pittsburgh, Pittsburgh, PA 15260, USA;

Anupam Banerjee − Laufer Center for Physical and Quantitative Biology, Stony Brook University, NY 11794, US; Department of Biochemistry and Cell Biology, Renaissance School of Medicine, Stony Brook University, NY 11794, USA;

Satyaki Saha − Laufer Center for Physical and Quantitative Biology, Stony Brook University, NY 11794, US; Department of Biochemistry and Cell Biology, Renaissance School of Medicine, Stony Brook University, NY 11794, USA;

Michael Ohlmeyer – Atux Iskay LLC, Plainsboro, New Jersey, NJ, 08536, USA;

Reuven Gordon − Department of Electrical and Computer Engineering, University of Victoria, Victoria, BC V8P 5C2, Canada;

Laura S. Itzhaki – Department of Pharmacology, University of Cambridge, Tennis Court Road, Cambridge CB2 1PD, UK;

Ivet Bahar − Laufer Center for Physical and Quantitative Biology, Stony Brook University, NY 11794, US; Department of Biochemistry and Cell Biology, Renaissance School of Medicine, Stony Brook University, NY 11794, USA;

## Author Contributions

Conceptualization: M.G. and I.B. Methodology: S.Z.Y., A.B., S.S., I.B., and M.G. Investigation: S.Z.Y., A.B., S.S., I.B., and M.G. Visualization: S.Z.Y., A.B., S.S., and M.G. Supervision: M.G. and I.B. Writing—original draft: S.Z.Y., A.B., S.S., R.G., L.S.I., I.B., and M.G. Writing—review and editing: S.Z.Y., A.B., S.S., M.O., R.G., L.S.I., I.B., and M.G.

## ACKNOWLEDGMENT

I.B. gratefully acknowledges support from the National Institutes of Health awards R01GM138287 and R01DK116780.

## ABBREVIATIONS

AMBER: Assisted model building with energy refinement
Cα: Alpha carbon
CHARMM: Chemistry at Harvard macromolecular mechanics
CoM: Center of mass
Cryo-EM: Cryo-electron microscopy
ENM: elastic network model
HEAT: Huntingtin Elongation factor 3–PP2A–TOR1
MEK: MAPK/ERK kinase
MD: Molecular dynamics
mTOR: Mammalian target of rapamycin
NAMD: Nanoscale molecular dynamics
PCA: Principal component analysis
PDB: Protein data bank
PP2A: protein phosphatase 2A
PRODIGY-LIG: Protein binding energy prediction for ligands
RMSD: Root-mean-square deviation
RMSF: Root-mean-square fluctuation
ROSIE: Rosetta online server that includes everyone
SMAP: Small molecule activator of PP2A
SMD: Steered molecular dynamics
TR: Tandem repeat
VMD: Visual molecular dynamics

