## Supporting Information for "Small molecule activator of phosphatase PP2A remodels scaffold PR65 structural dynamics to promote holoenzyme assembly"

### Supplementary Methods

**Molecular dynamics (MD) simulation details for ATUX-8385 bound at S3.** The PR65-SMAP complex with ATUX-8385 bound at site S3 was solvated within a truncated octahedral box using the TIP3P water model[1], maintaining a minimum distance of 26.0 Å between the solute and the box edges. Sodium ($Na^+$) and chloride ($Cl^-$) ions were added to achieve charge neutrality. All preparatory steps were carried out using the *tLEaP* module from AmberTools. Three independent MD simulations were performed using the AMBER20 software[2]. The ff14SB force field[3] was applied to the protein and the general Amber force field (GAFF)[4] was employed for small molecules. Ligand charges were derived from AM1-BCC calculations[5] using *antechamber*[6] and *parmchk2* tools. Each system underwent a multistep minimization and equilibration protocol. Initially, the systems were subjected to 2000 steps of unrestrained minimization, using the steepest descent method for the first 500 steps to manage high-energy configurations, followed by 1500 steps of conjugate gradient minimization. Subsequently, the systems underwent a 20 ps restrained equilibration under the NVT ensemble at 298 K, employing a Langevin thermostat with a damping coefficient of 1 $ps^{-1}$ and harmonic restraints (force constant of $k$ = 1 kcal/mol/$Å^2$) applied to all non-hydrogen atoms except those in water and ions. This was followed by a 1 ns restrained equilibration under NPT conditions, maintaining pressure at 1 atm using the Monte Carlo (MC) barostat, while retaining harmonic restraints. Subsequently, harmonic restraints were removed, and an unrestrained NPT equilibration phase of 1 ns was conducted. After this final equilibration phase, production MD simulations of 800 ns length were performed for each system. A 2 fs timestep was used in all simulation's phases. Long-range electrostatic interactions were computed using the particle-mesh Ewald (PME) method, and non-bonded interactions were truncated at a 10 Å cut-off.

### Supplementary Figures

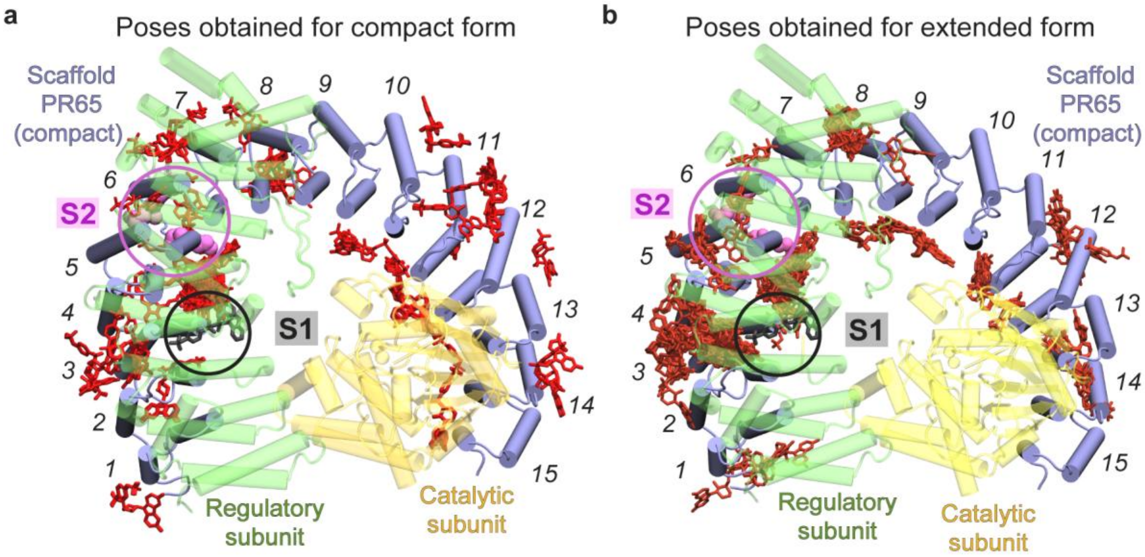


**Figure S1.** Binding poses from blind docking simulations superimposed onto the compact PR65 of trimeric PP2A structure (PDB: 6NTS[7]). The observed binding sites of ATUX-8385 (*red sticks*) onto PR65 (*blue cartoon*) in the (a) compact (PDB: 6NTS[7]) and (b) extended (PDB: 1B3U[8]) forms are depicted on the trimeric PP2A for visualization. Binding poses were obtained for PR65 alone and are shown on the PP2A holoenzyme (PDB: 6NTS[7]) by superimposing the PR65 structures. For panel (a), no alignment was necessary since the PR65 structure was taken directly from the trimer; for panel (b), the extended PR65 was superimposed onto the compact PR65 by aligning HEAT repeats 1–9. DT-061 resolved by cryo-EM (PDB: 6NTS[7]) is shown in *black sticks* (inside a *black circle*) to indicate its binding site, S1. The *magenta spheres* (inside the *magenta circles*) refer to three residues, K193, E196, and L197, proposed to be involved in binding a SMAP to the PR65 scaffold. The catalytic and regulatory subunits (not included in docking simulations) are shown as *yellow* and *green*, respectively.

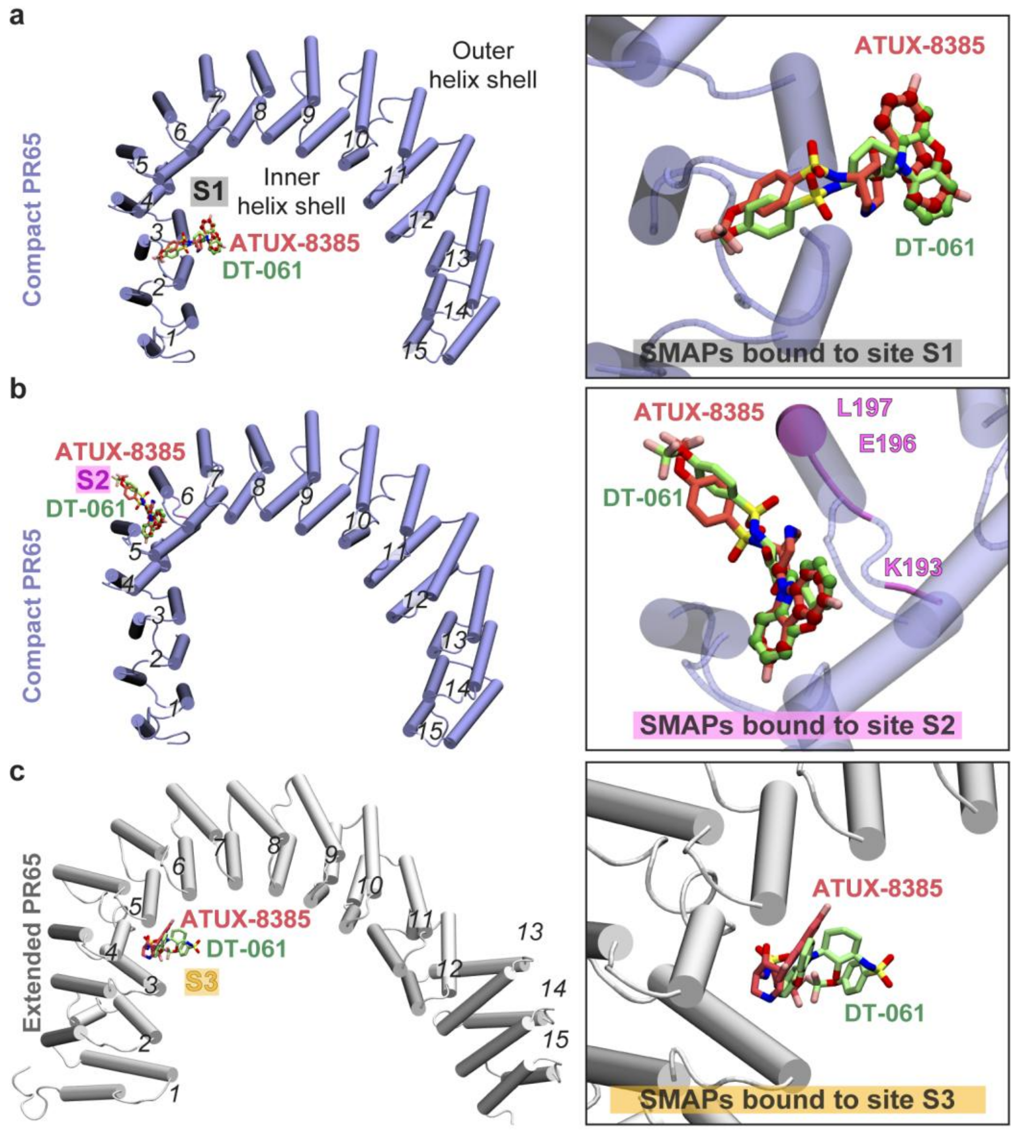


**Figure S2.** SMAP binding poses used as starting conformations for MD simulations. *In silico* binding of ATUX-8385 (*red*) and DT-061 (*green*) to (a) S1, (b) S2, and (c) S3 of PR65. (a) DT-061 bound at S1 in the compact PR65 conformation, as observed in the cryo-EM resolved PP2A complex (PDB: 6NTS[7]). ATUX-8385 was modeled at S1 by aligning its carbon atoms (C11-C17, C19-C22; *red beads*) to those of DT-061 (C1-C3, C6-C12, C18; *green beads*). (b) Docked pose of ATUX-8385 bound at S2 of PR65 in the compact conformation near the K193, E196, and L197 residues *(*highlighted in *magenta*), a region previously[9] experimentally implicated in coordinating SMAP interactions with PR65. The pose was obtained using the Rosetta Ligand Docking protocol[10] via the ROSIE server[11], with the experimentally identified site K193-L197 used as a constraint. For modeling DT-061 binding at the S2 site, we then reversed this alignment procedure, aligning DT-061's carbon atoms (C1–C3, C6–C12, C18) to those of ATUX-8385 (C11–C17, C19–C22) at S2. (c) Docked poses of ATUX-8385 and DT-061 bound to PR65 at S3 in the extended conformation, generated using Autodock Vina[12] with the K193–L197 site as a reference.

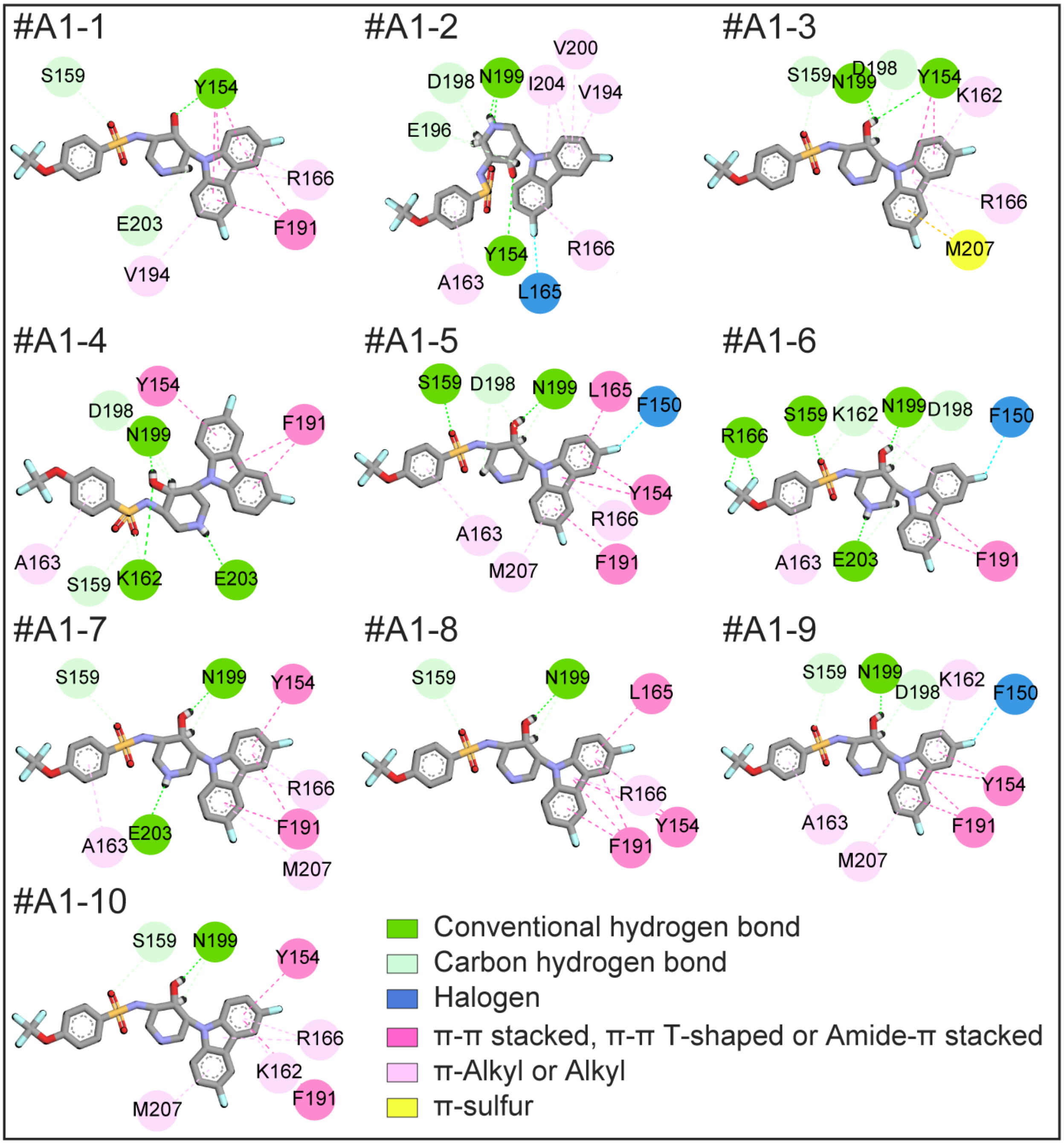


**Figure S3.** Interactions of 10 randomly selected ATUX-8385 conformations bound to S2 in binding pose S2-A1. Interactions between ATUX-8385 and PR65 are shown for 10 randomly selected ATUX-8385 conformations in binding pose S2-A1. Discovery Studio Visualizer[13] was used to determine and visualize interactions.

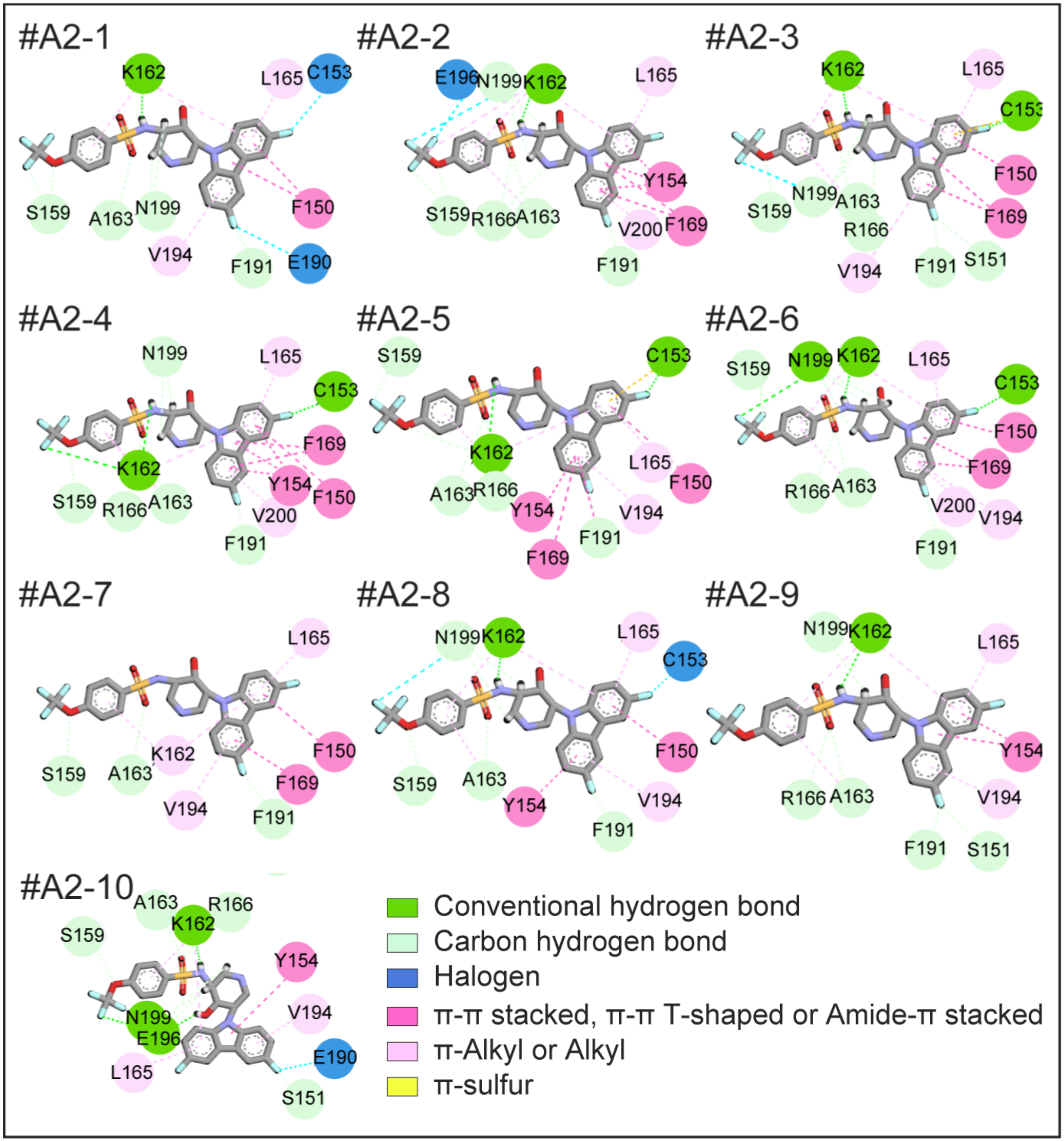


**Figure S4.** Interactions of 10 randomly selected ATUX-8385 conformations bound to S2 in binding pose S2-A2. Interactions between ATUX-8385 and PR65 are shown for 10 randomly selected ATUX-8385 conformations in binding pose S2-A2. Discovery Studio Visualizer[13] was used to determine and visualize interactions.

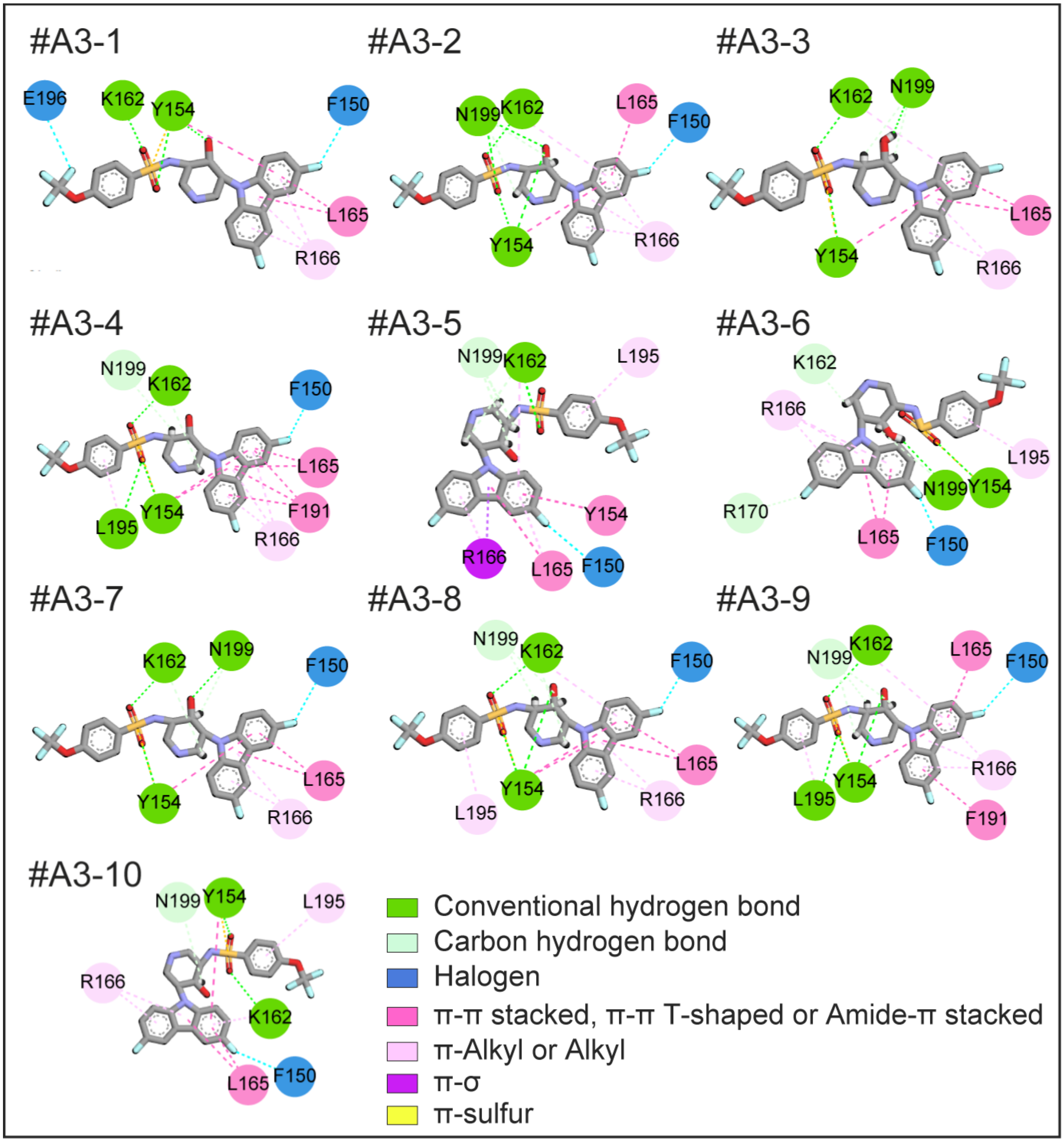


**Figure S5.** Interactions of 10 randomly selected ATUX-8385 conformations bound to S2 in binding pose S2-A3. Interactions between ATUX-8385 and PR65 are shown for 10 randomly selected ATUX-8385 conformations in binding pose S2-A3. Discovery Studio Visualizer[13] was used to determine and visualize interactions.

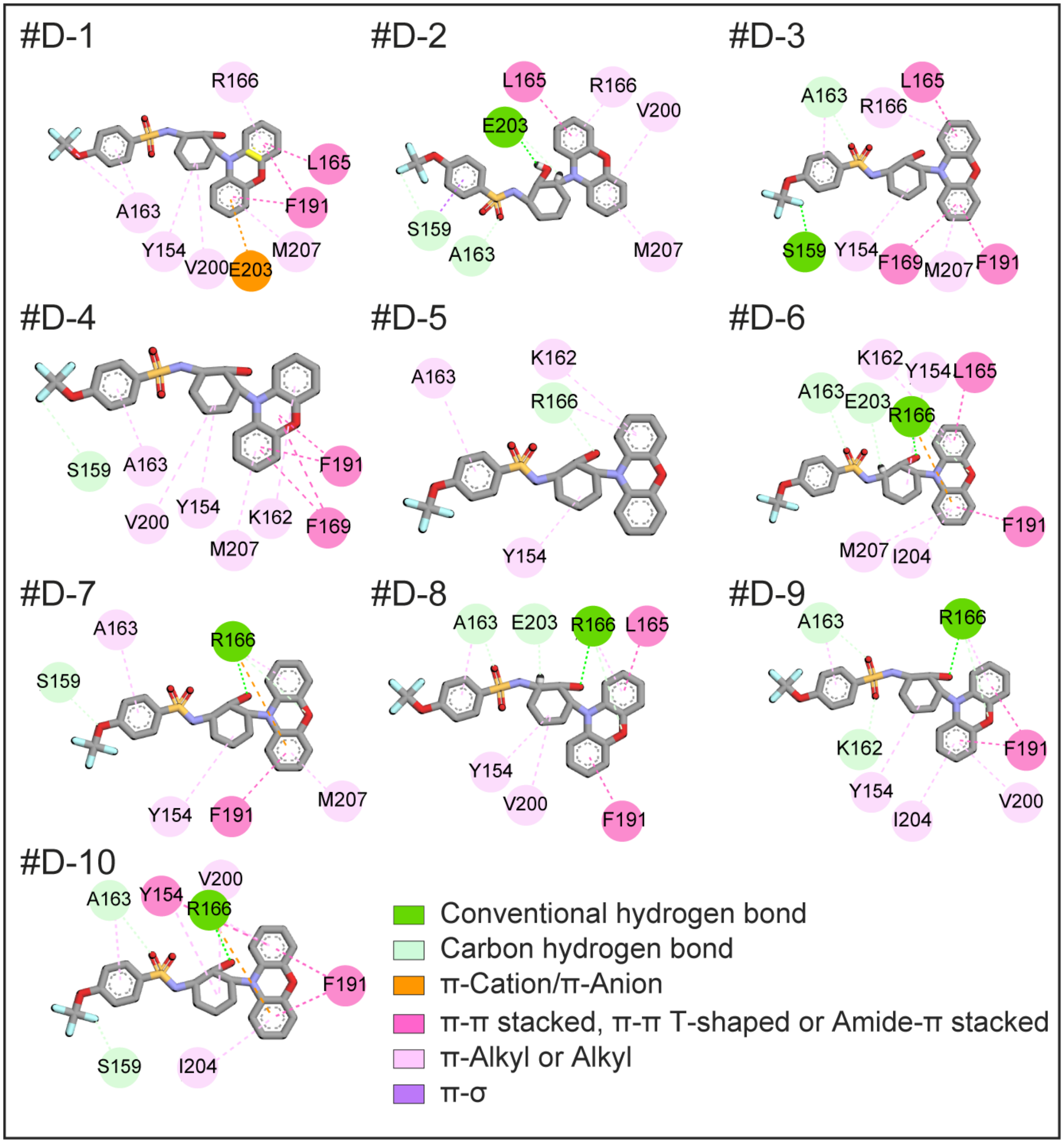


**Figure S6.** Interactions of 10 randomly selected DT-061 conformations bound at S2 in binding pose S2-D. Interactions between DT-061 and PR65 are shown for 10 randomly selected DT-061 conformations in binding pose S2-D. Discovery Studio Visualizer[13] was used to determine and visualize interactions.

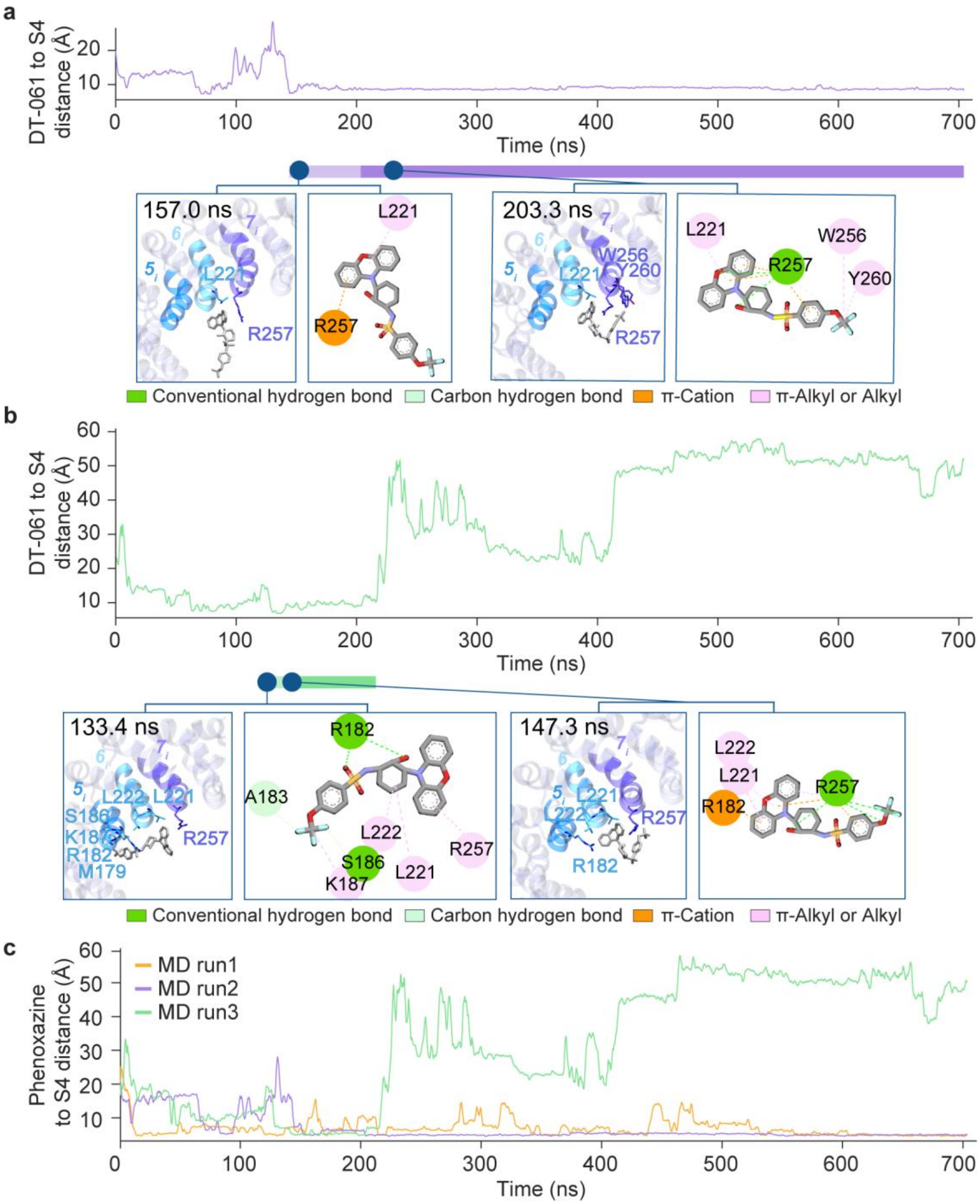


**Figure S7.** Time evolution of the DT-061 binding pose during MD simulations, and interactions stabilizing the binding of the SMAP. (a-b) The distances between the center of mass (CoM) of DT-061 and the CoM of the N-terminal halves of helices $5_i$-$7_i$ (P178-A183, D217-L222, and R257-M261) are plotted as a function of time, for the two runs initiated with DT-061 originally bound to S1 (see the third in **Figure 4**). Horizontal bars between the time evolution curves indicate the trajectory stretches used for PCA. The *darker portions* of the bars represent the parts of the trajectory during which the stable state S4 was predominantly sampled. Each binding conformation is depicted through a diagram generated using VMD[14] (*left*), and a schematic illustration, from the Discovery Studio Visualizer[13], which showcases the atomic interactions between DT-061 and PR65 (*right*). (c) The evolution of distances between the CoM of DT-061 phenoxazine and the CoM of the N-terminal halves of helices $5_i$-$7_i$.

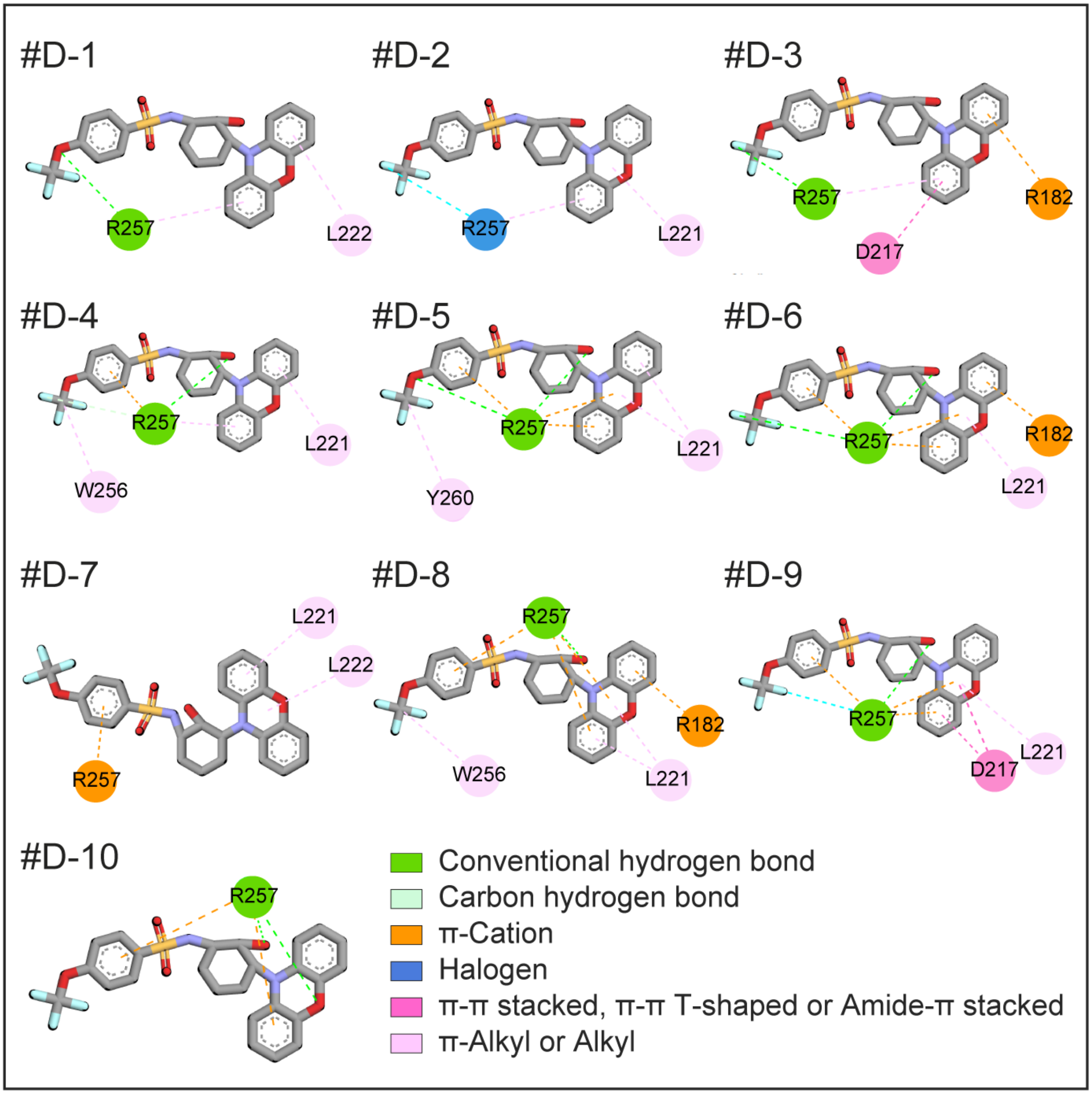


**Figure S8.** Interactions of 10 randomly selected DT-061 conformations bound to S4. Interactions between DT-061 and PR65 are shown for 10 randomly selected DT-061 conformations in pose S4. Discovery Studio Visualizer[13] was used to determine and visualize interactions.

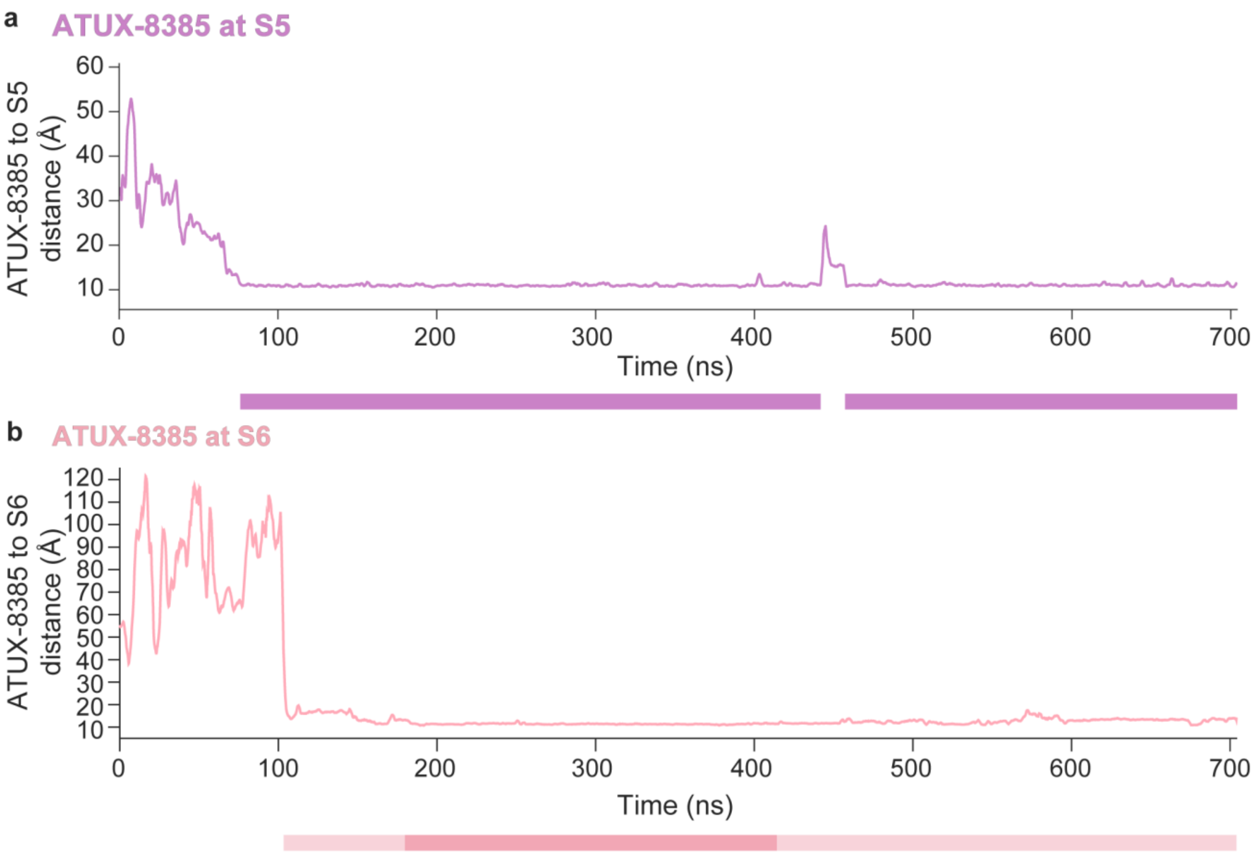


**Figure S9.** Time evolution of the distance of ATUX-8385 to S5 and S6 binding sites observed in MD simulations. (a) The evolution of distances between the CoM of ATUX-8385 and the CoM of the C-terminal halves of segments $8_o$ and $6_i$-$8_i$ (P275-N288, L222-Q232, M261-V273, and A300-N311) in set V run with ATUX-8385 at S5. (b) The evolution of distances between the CoM of ATUX-8385 and the CoM of segments $12_o$, and $12_i$-$14_i$ (V435-V451, Y455-F472, Y494-C511, A533-I549) in set V with ATUX-8385 at S6. Bars under the plots indicate the trajectory stretches that were used for PCA, with the *darker portions* corresponding to the minima in the subspace of principal components presented in **Figure S10**.

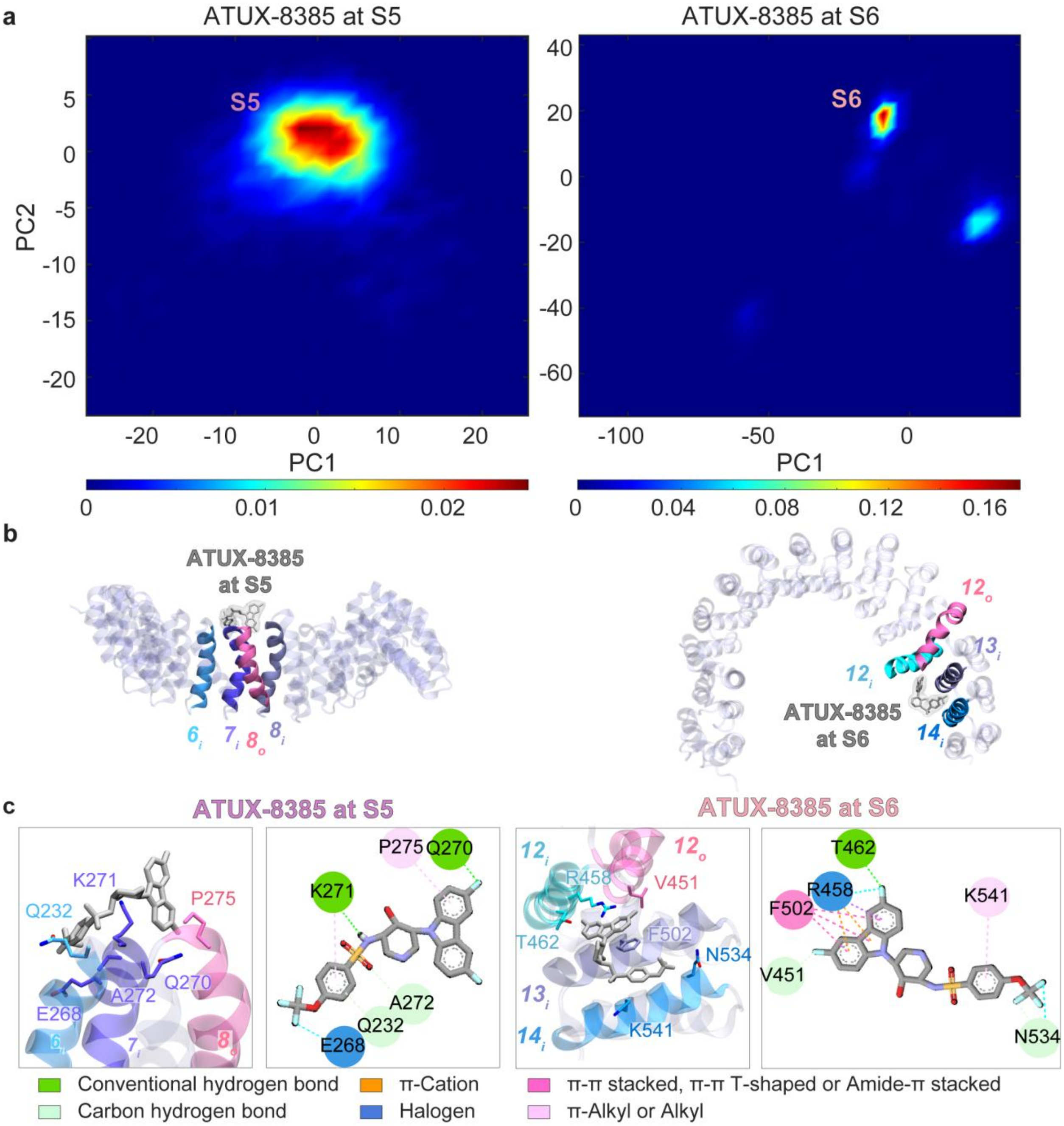


**Figure S10.** ATUX-8385 binding poses observed in MD simulations. (a) The distributions of SMAP binding poses, projected onto the principal space of bound conformers extracted from MD trajectories for ATUX-8385 at S5 (*left*) and S6 (*right*). Conformational maps are shown for ATUX-8385 bound to S5, with one leading binding pose S5, and ATUX-8385 bound to S6, with a single dominant binding pose S6. (b) The most frequently observed SMAP binding poses, as identified in panel (a), are shown. (c) Each binding pose is depicted through two distinct representations as described in previous figures. In site S5, ATUX-8385 interacts with $6_i$ Q232, with $7_i$ E268, Q270, K271, and A272, with $8_o$ P275, and $8_i$ N311. The most frequent interactions are those with E268, Q270, K271, A272, and P275. When bound to site S6, ATUX-8385 is coordinated by $12_o$ V451, $12_i$ R458 and T462, $13_i$ Y494, M498, and F502, and $14_i$ N534, F537, N538, K541, and K545. The most frequent interactions were those with V451, T462, R458, F502, N534, F537, and K541.

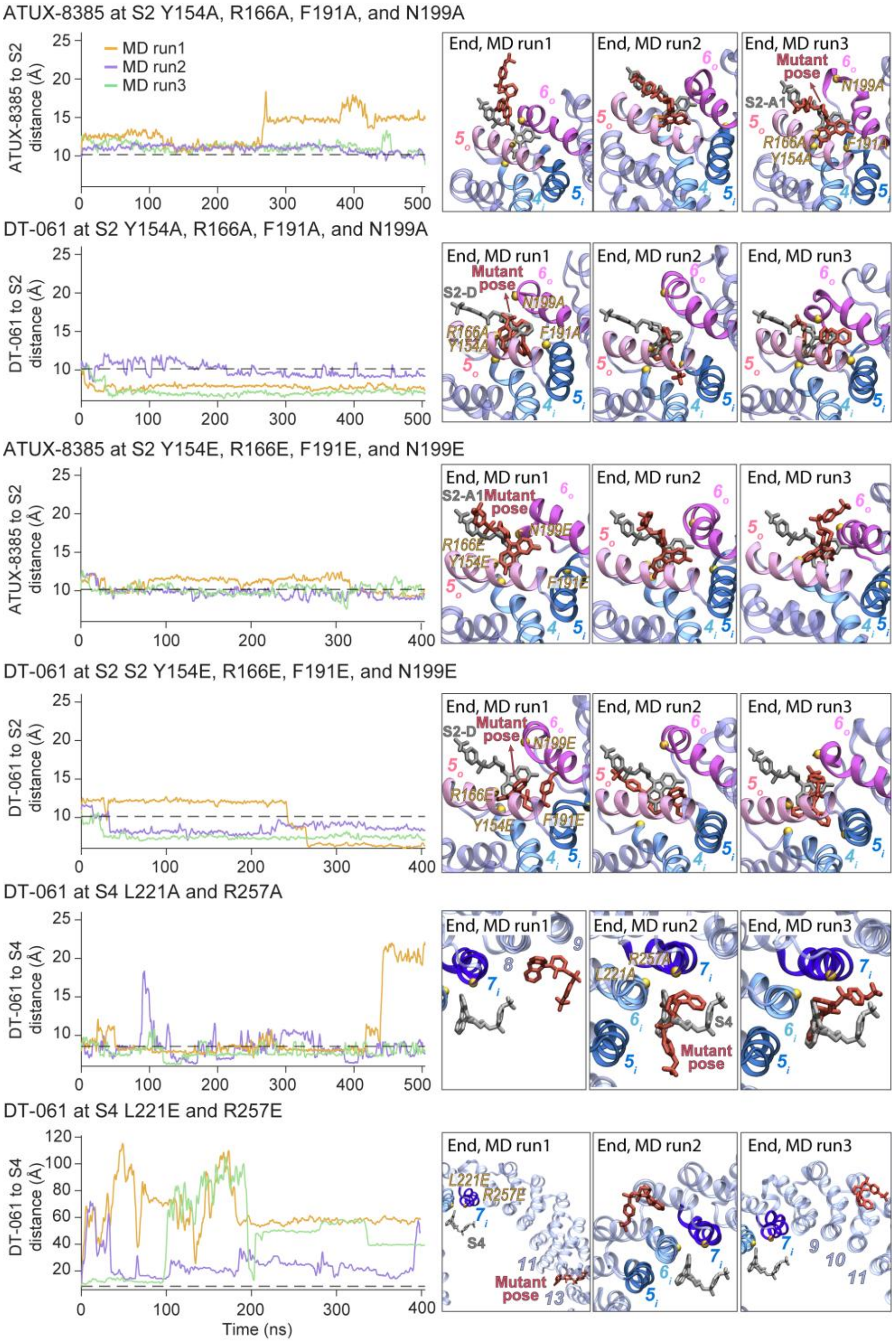


**Figure S11.** Time evolution of SMAP–binding site distances in mutant MD simulations and endpoint poses. *Left panels:* For simulations initiated with SMAPs bound at S2, the distance of each SMAP to the S2 site was calculated as the separation between the CoM of each SMAP and the CoM of PR65 helices $4_i$, $5_i$, $5_o$, and $6_o$ (residues F140–Y154, A160–S174, P178–A192, and L197–S213, respectively). For simulation initiated with SMAPs at S4, the distance between the SMAP and S4 was calculated using the CoM of each SMAP and the CoM of the N-terminal halves of PR65 helices $5_i$–$7_i$. Each trace (*orange*, *purple*, and *green*; run1-3, respectively) represents one of three independent MD runs starting from the same binding pose and PR65 mutant. *Right panels:* Endpoint snapshots showing the final bound pose captured at the last frame of each trajectory for ATUX-8385 or DT-061 in the indicated mutant backgrounds.

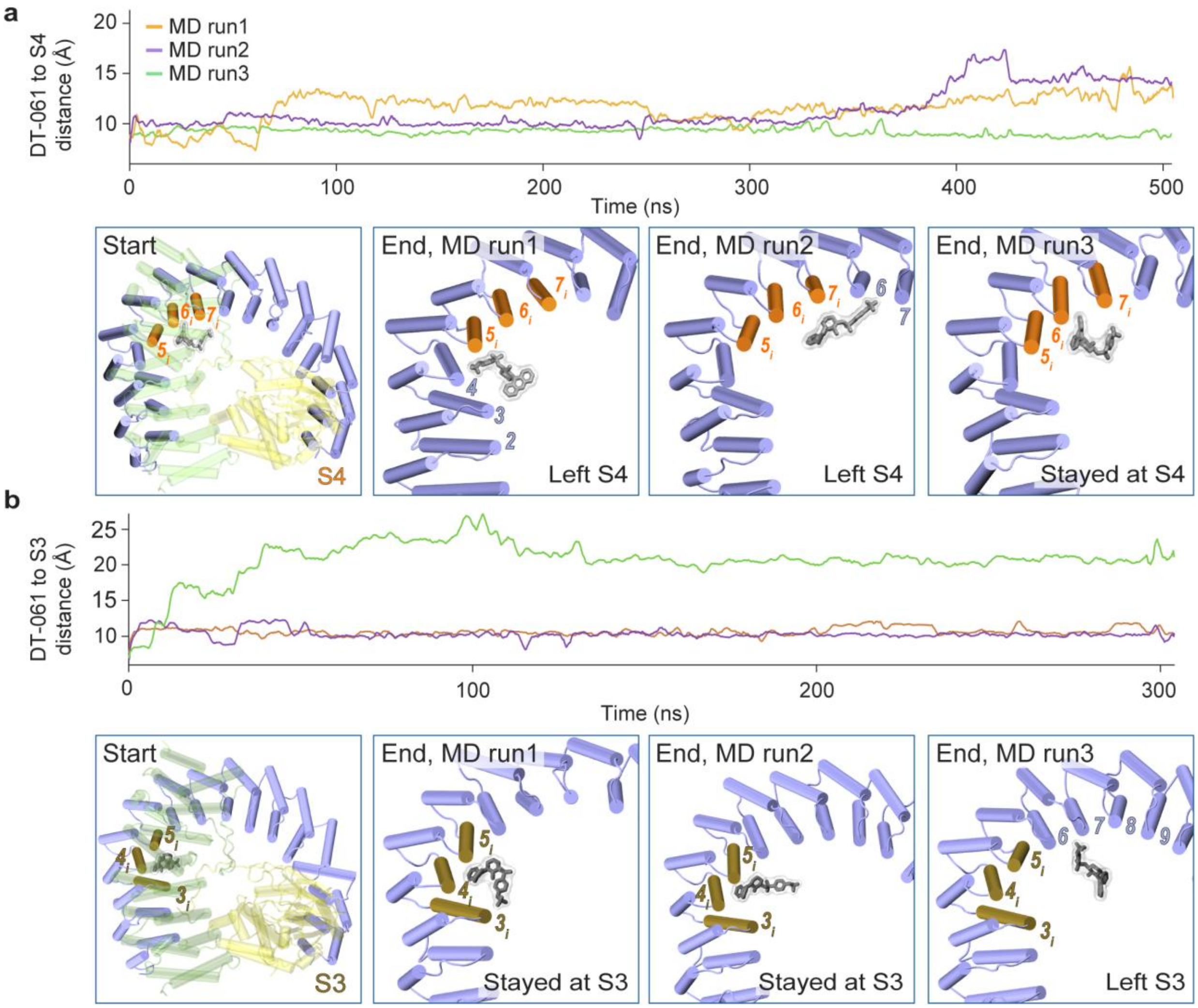


**Figure S12.** DT-061 seeded at S4 (a) or S3 (b) in MD simulations of the trimeric PP2A. (a) DT-061 to S4 distance is the CoM separation between DT-061 and the CoM of the N-terminal halves of PR65 helices $5_i$–$7_i$. (b) DT-061 to S3 distance is the CoM separation between DT-061 and the CoM of PR65 helices $3_i$–$5_i$ (residues T101–E117, F140–Y154, and P178–Q192). Distances are plotted versus time for three independent trajectories per condition (~500 ns in panel (a); ~300 ns in panel (b)). Snapshots below each plot show the starting configuration (*left panel*) and the end-of-trajectory poses for the three independent runs (*middle and right panels*). Although all simulations were performed with the full trimeric complex, only PR65 is shown in the end-state panels for clarity. Annotations indicate whether DT-061 remained in the starting pocket or left it. All simulations were performed with the full trimeric complex, but only PR65 is shown in the end-state panels for clarity. PR65 is shown in *blue*; the catalytic and regulatory subunits are *yellow* and *green*, respectively; and DT-061 is shown in *gray*.

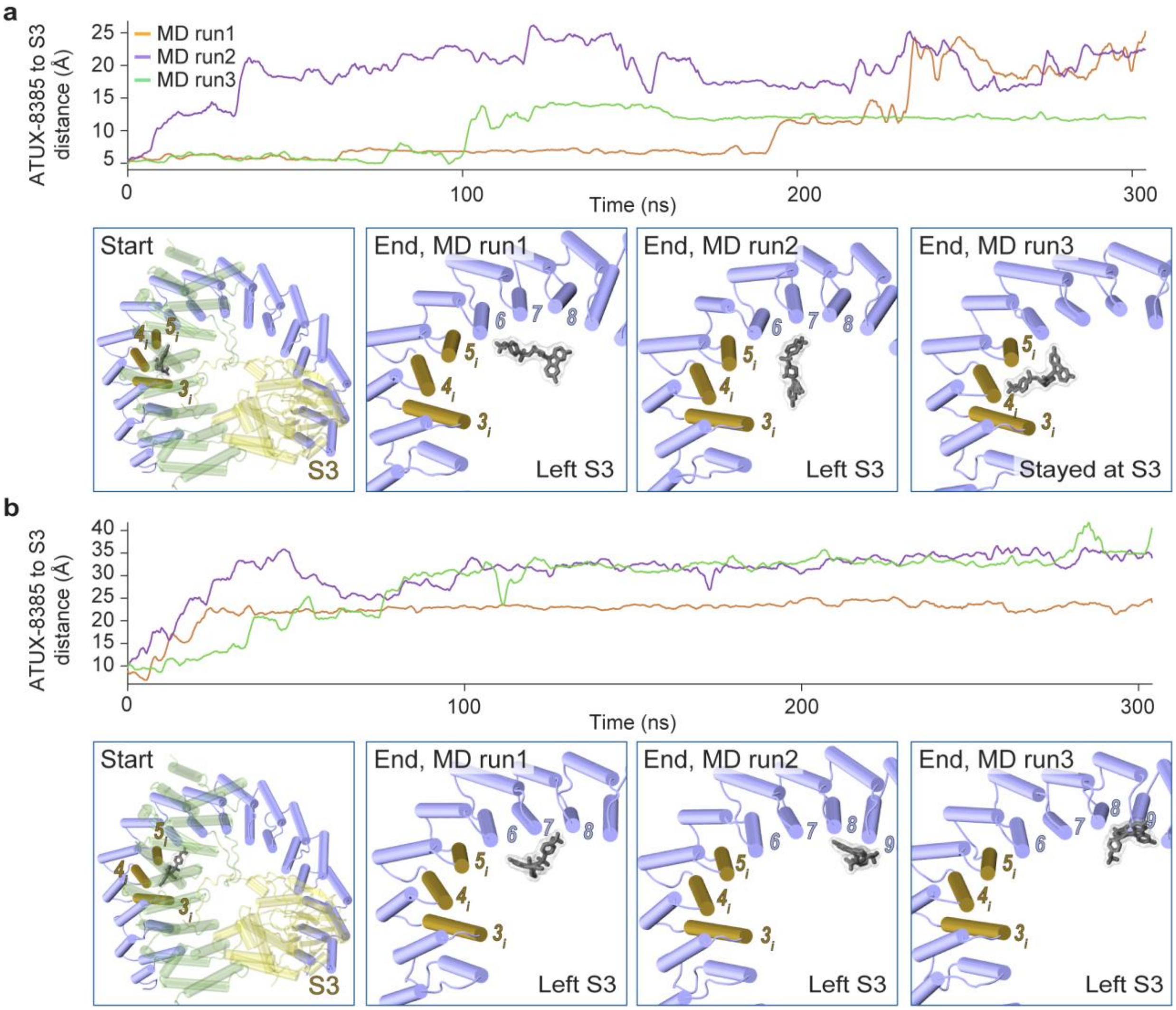


**Figure S13.** ATUX-8385 seeded in the S3 of trimeric PP2A: distance traces and endpoint snapshots. Trimeric PP2A simulations were initiated from S3 (a) poses obtained by docking ATUX-8385 to monomeric PR65 (in extended form) and (b) poses derived from an MD-refined S3 bound conformation. ATUX-8385 to S3 distance is the separation between ATUX-8385 CoM and the PR65 S3 (helices $3_i$–$5_i$) CoM. Distances are shown over simulation time for three independent trajectories per starting pose (~300 ns each). Snapshots beneath each plot show the starting configuration and the final frame of each trajectory. Labels indicate whether the ligand remained in S3 or dissociated. All simulations were performed with the full trimeric PP2A holoenzyme; for clarity, only PR65 is shown in the simulation endpoint panels. PR65 is colored *blue*, the regulatory subunit *green*, the catalytic subunit *yellow*, and ATUX-8385 *gray*.

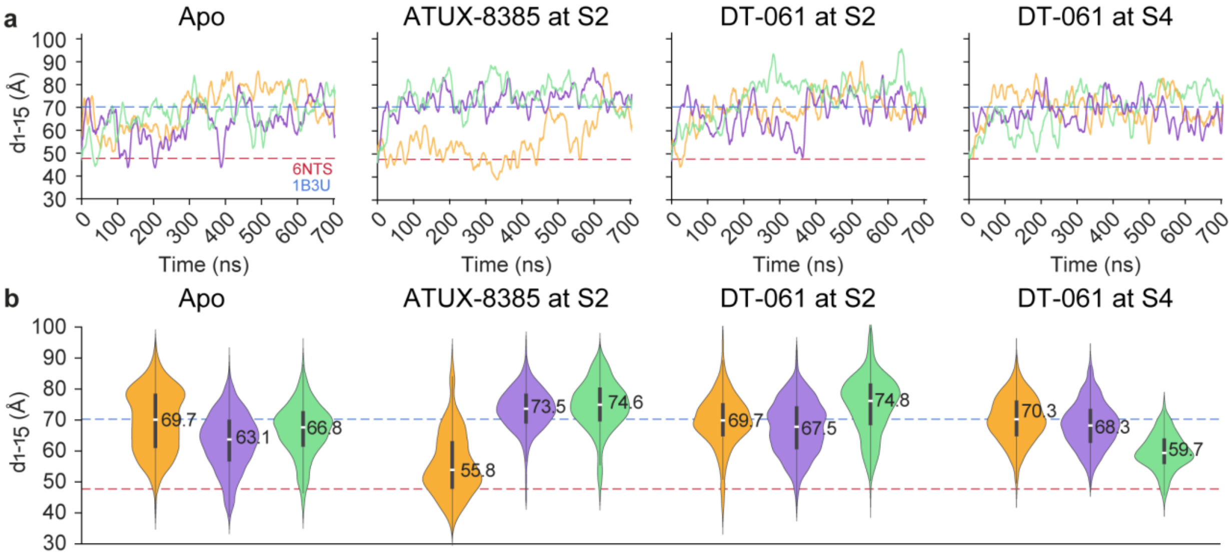


c

| | Apo | ATUX-8385 at S2 | DT-061 at S2 | DT-061 at S4 |
|---|---|---|---|---|
| PC1 | 0.76 | 0.60 | 0.74 | 0.73 |
| PC2 | 0.41 | 0.58 | 0.48 | 0.38 |
| PC3 | 0.10 | 0.41 | 0.30 | 0.32 |
| PC4 | 0.13 | 0.03 | 0.22 | 0.28 |
| PC1-2 | 0.86 | 0.83 | 0.88 | 0.82 |

**Figure S14.** Distribution of end-to-end distances in apo and SMAP-bound PR65 from MD simulations. (a) Time evolution of the end-to-end distances ($d_{1\text{-}15}$). The contributions of runs 1-3 in each case are shown in *orange*, *purple*, and *green*, respectively. *Dashed lines* depict the end-to-end distances observed in the compact (PDB: 6NTS[7]) and extended (PDB: 1B3U[8]) structures of PR65. (b) Violin plots representing the distribution of end-to-end distances for individual runs. (c) Correlation cosines between the PCs sampled in simulations and the deformation vector experimentally observed between the compact and extended forms of PR65. The correlation cosines refer to that between (i) the four most significant PCs of the apo and SMAP-bound PR65 derived from the combined (triplicate) trajectories, and (ii) the deformation vector that delineates the transition from compact to extended PR65 structures. The y-axis refers to PC1-PC4, individually, and the last row shows the cumulative effects of PC1-PC2. The correlation cosines are color-coded: *white* signifies no correlation, while *blue* indicates a complete correlation.

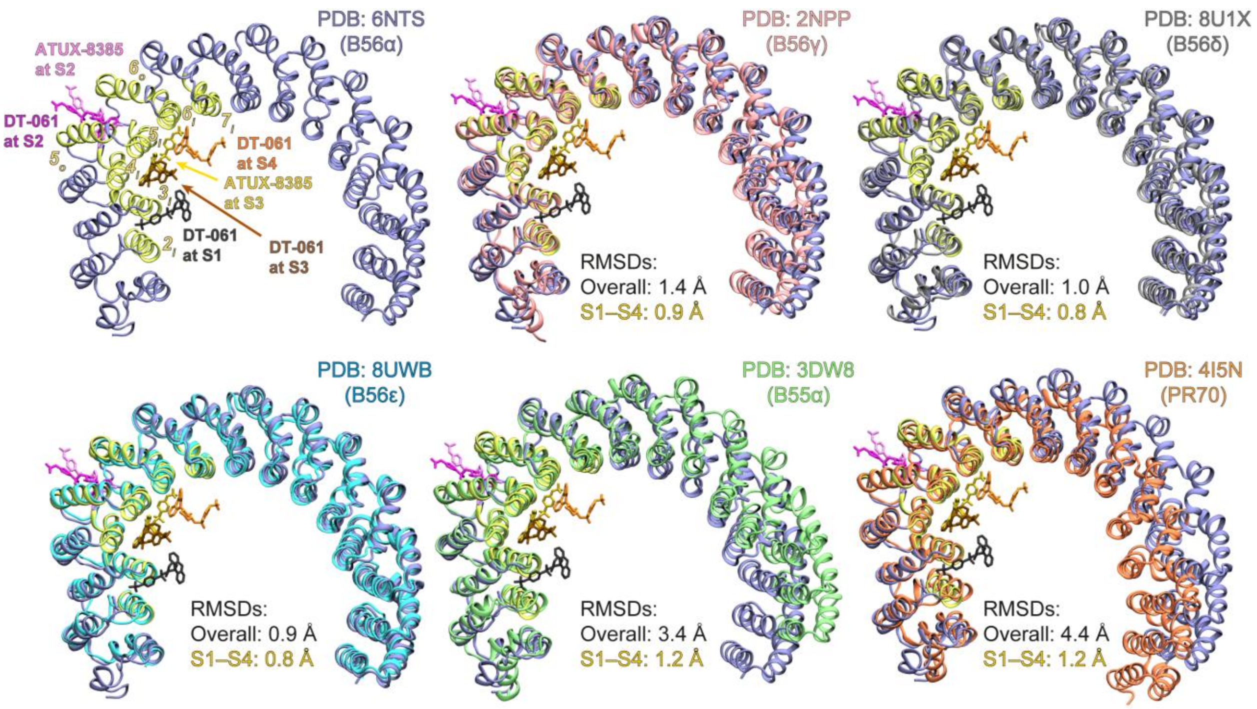


**Figure S15.** Superposition of PP2A heterotrimers containing distinct regulatory B family members: B56α (PDB: 6NTS[7]; reference), B56γ (PDB: 2NPP[15]), B56δ (PDB: 8U1X[16]), B56ε (PDB: 8UWB[17]), B55α (PDB: 3DW8[18]), and PR70 (PDB: 4I5N[19]). In the reference trimeric complex, only the PR65 scaffold is shown in *iceblue*, and the extended SMAP-binding region (sites S1–S4) is highlighted in *yellow*. The PR65 subunits of PP2A holoenzymes with the other B subunits were superposed using two protocols: (i) S1–S4 helices only, which is shown here, and (ii) all PR65 helices. Helix-only RMSDs for S1–S4 and for the entire PR65, each computed relative to reference (PDB:6NTS[7]), are reported in the panels. Representative SMAP poses at sites S1–S4, derived from the cryo-EM resolved DT-061 pose at S1, the docked DT-061 pose at S3, and MD-refined conformations of DT-061 and ATUX-8385 at S2-S4, are shown in *licorice representation* and were superimposed on each structure depicted in this figure.
